# NAC promotes co-translational protein folding at the ribosomal tunnel exit

**DOI:** 10.1101/2025.08.02.668148

**Authors:** Jaime Santos, Manuel Günnigmann, Radoslaw J. Gora, Marija Iljina, Masa Predin, Ilgin Eser Kotan, Pratiman De, Dhawal Choudhary, Juwon Jang, Frank Tippmann, Nenad Ban, Sander J Tans, Shu-ou Shan, Günter Kramer, Bernd Bukau

**Author notes:** These authors contributed equally (Jaime Santos, Manuel Günnigmann, Radoslaw J. Gora). **Lead contact** Further information and requests for resources and reagents should be directed to and will be fulfilled by the lead contact, Bernd Bukau. Correspondence (B.B.), (G.K.), (S.S.), (S.J.T.), (N.B.).

## Abstract

The nascent polypeptide-associated complex (NAC) coordinates enzymatic modifications and membrane targeting of nascent chains during translation. While NAC’s function as a dynamic hub for other factors is well-established, its direct role in co-translational folding is unclear. By proteome-wide profiling NAC co-translational interactions in human cells, we found that NAC recognizes emerging segments enriched in hydrophobicity and α-helical propensity, within folded domains of cytonuclear proteins. Single-molecule and structural analyses reveal that NAC, via its β-barrel domain, dynamically interacts with nascent chains at the ribosomal tunnel exit and is capable of promoting on-pathway folding. Compartment-specific nascent chain interactions of NAC further elucidate its role in targeting to the endoplasmic reticulum and mitochondrial membrane protein biogenesis. Together, these findings show that NAC acts as a bona fide co-translational chaperone that facilitates early protein folding at the ribosomal tunnel exit, expanding its functional repertoire in protein biogenesis.

## Introduction

To achieve biological activity, newly synthesized proteins must adopt their native structures, undergo N-terminal processing, and be targeted to specific cellular compartments. These processes begin co-translationally, mediated by numerous factors that interact with the growing polypeptide chains^1–10^. The nascent polypeptide-associated complex (NAC) is unique among these factors. By binding near the ribosomal polypeptide exit tunnel^3,11,12^, NAC serves as a coordinating hub, recruiting other factors to the ribosome and regulating their access to nascent chains (NC)^2–4,13–15^. Hence, NAC is implicated in various co-translational processes, including NC targeting to the endoplasmic reticulum (ER) and mitochondria and coordination of enzymatic modifications such as methionine excision and N-terminal acetylation^2–4,12,13,16^.

NAC is a highly conserved heterodimeric complex comprising NACα and NACβ (also known as BTF3 in humans). Its binding to the ribosome is primarily mediated by a positively charged anchor motif in the N-terminus of NACβ (**Figure 1A**)^17,18^. The central β-barrel domain, formed by both subunits (**Figure 1A**), interacts with the ribosome near the polypeptide tunnel exit. This interaction is dynamic and can be displaced, allowing NCs to become accessible to other biogenesis factors^3,4^. The C-terminal tails of NACα and NACβ help recruit various factors, including the signal recognition particle (SRP), methionine amino peptidase (MetAP1), N-acetyl transferase A (NatA), and the eEF1A-specific chaperone Chp1 (**Figure 1A**)^3,4,13,14^, establishing NAĆs role as hub for NC-interacting factors and suggesting involvement in yet undiscovered co-translational processes.

**Figure 1:**
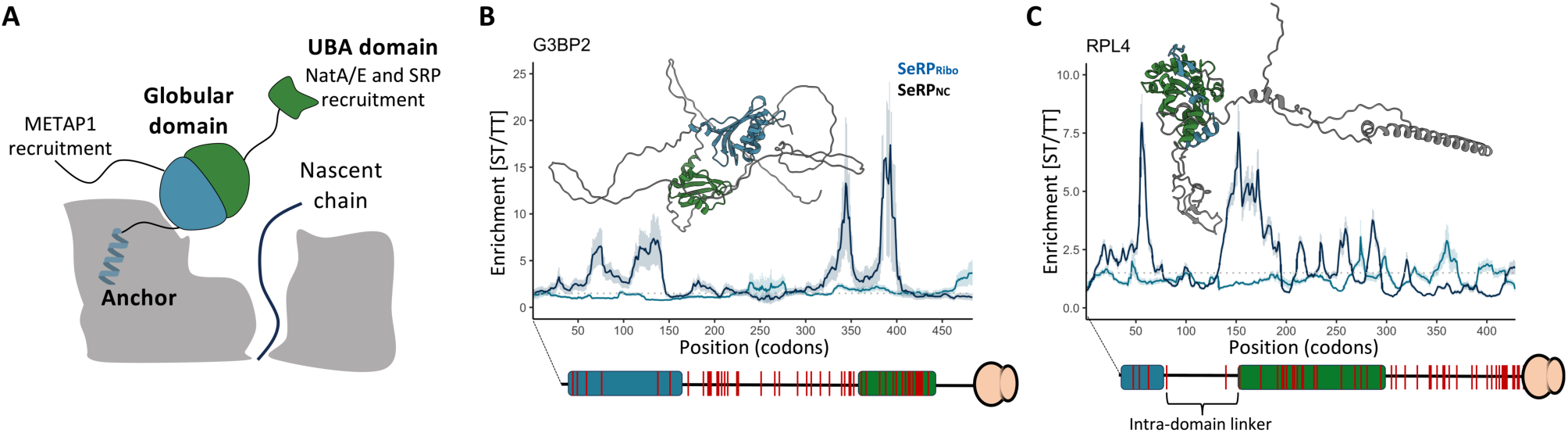
Structural features of NAC for development of a Selective Ribosome Profiling strategy. (**A**) Schematic representation of NAC association with the ribosome and domain architecture of NACα (green) and NACβ (blue). (**B-C**) Enrichment profiles of nascent cytonuclear proteins G3BP2 and RPL4 along coding sequences. NC independent (SeRP_Ribo_) and dependent (SeRP_NC_) enrichments are colored in blue and black, respectively. Protein domains (rectangles) and lysines (red bars) are annotated using a 30-residue tunnel correction to account for nascent chain emergence from the exit tunnel, as depicted in Figure S1A. Shown structures were extracted from AlphaFold^66,67^. The dotted line indicates an enrichment of 1.5, used as a threshold to define binding periods. Shadowed areas indicate 95% confidence interval.

In addition to its coordinating roles, NAC has also been proposed to function as a chaperone^19–23^, though its exact role in NC folding is unclear. NAC inhibits protein aggregation post-translationally^24–26^, but evidence for a chaperoning function at the ribosome is indirect. NAC depletion in human and yeast increases co-translational ubiquitination of NCs^20,21^, and simultaneous deletion of NAC and the Hsp70 homolog Ssb in yeast elevates protein aggregation^27^. Importantly, NAĆs post-translational chaperone function requires its ribosome anchor region, which is inaccessible for substrate interactions when associated with the ribosome (**Figure 1A**)^25^. Since NAC acts as a central coordinator of co-translational factors^2–4,14^, phenotypes initially linked to its proposed chaperone role may indirectly arise from the missing regulatory activity. Thus, the co-translational folding function of NAC remains a topic of debate.

Here, we characterize NAC’s proteome-wide interactions with NCs *in vivo* and investigate its proposed chaperoning functions *in vitro*. By developing NAC-selective ribosome profiling (SeRP) and integrating it with single-molecule Förster resonance energy transfer (smFRET), single particle cryo-electron microscopy (cryo-EM), and optical tweezer experiments, we describe the basis of NAC’s co-translational chaperoning activities. NAC’s β-barrel domain interacts with emerging polypeptides at the ribosomal tunnel exit, suggesting that it facilitates early NC folding events. Furthermore, NAC interactions facilitate on-pathway folding of a model protein, minimizing folding delays and preventing incomplete folding intermediates. Our SeRP analysis also reveals compartment-specific NC binding functions of NAC, corroborating its participation in targeting ER-translocated proteins, chaperoning cytonuclear and mitochondrial proteins, and assisting mitochondrial membrane protein biogenesis. Collectively, these findings provide a comprehensive perspective on NAC-mediated protein biogenesis, unveiling a novel type of chaperoning activity of this central ribosome-associated factor.

## Results

### Design of a NAC-selective ribosome profiling strategy

To investigate NAĆs role in protein biogenesis, we developed a SeRP strategy to analyze its interactions with the nascent proteome of human HEK293-T cells. SeRP enables codon-resolution mapping of NAC interactions during *in vivo* translation, facilitating the study of co-translational chaperone engagement and protein assembly^8,10,28–34^. This method involves the deep sequencing of mRNA fragments protected by the ribosome during decoding, also termed footprints. Specifically, we compare the total translatome (TT), which are the mRNA footprints for all ribosomes, to a selected translatome (ST), which comprises the footprints of immunopurified NAC-bound ribosomes. The enrichment of ST over TT indicates NAC binding to translating ribosomes across the pronome (**Figure S1A)**^31,34^.

NAC binds to all translating ribosomes with low nanomolar K_d_ (∼1 nM) via its anchor motif (**Figure 1A**) regardless of NC nature^3,17,18,35,36^. The outlined SeRP method will therefore capture NC-independent interactions (SeRP_ribo_), providing a comprehensive view of NAĆs ribosome association.

To understand NAC binding to NCs, we employed an alternative strategy. In SeRP, crosslinking approaches are often employed to stabilize chaperone-RNC complexes^30,34,37^. Using the lysine-specific homo-bifunctional crosslinker DSP (Dithiobis(succinimidylpropionate)), we found that NAC preferentially crosslinks to NCs of varying lengths instead of ribosomal proteins, as indicated by a smear rather than discrete bands in non-reducing western blots (**Figure S1B**). Notably, the lysines in the ribosome docking site of NAC contact ribosomal RNA instead of protein, and are therefore not expected to generate NAC-ribosome crosslinks^3^ (**Figure S1C**). To enrich crosslinked NAC-NC complexes, we took advantage of the fact that NAC dissociates from ribosomes during sucrose cushion and gradient fractionation (**Figures S1D and S1E**). *In vivo* crosslinking prior to this fractionation yielded a 3-fold increase in NAC co-sedimentation with ribosomes, indicating stabilized NAC-NC complexes (**Figure S1E**). This approach, termed SeRP_NC_, thus captures NAC interactions with NCs.

We tested both SeRP strategies in HEK293-T cells (**Figure S1F**) by analyzing five cytonuclear proteins (**Figures 1B, 1C and S1G-S1I**). For SeRP_ribo_, we observed flat binding profiles with enrichment values close to one across coding sequences (CDS), indicating uniform ribosome occupancy consistent with NC-independent binding. In contrast, SeRP_NC_ revealed distinct enrichment peaks, suggesting that NC features promote interaction with NAC at specific points of translation. To ensure that the SeRP_NC_ data were not biased by lysine distribution within the NCs, we compared NAC enrichments with lysine distribution and found no correlation (**Figures 1B, 1C and S1G-S1I**). For example, regions with high lysine density, such as the interdomain linker of G3BP2 and the C-terminal tail of RPL4 (red bars in **Figures 1B and 1C**, respectively), showed no significant NAC binding. These results indicate that DSP crosslinking effectively stabilized NAC interactions with NCs without introducing bias.

### NAC binding varies across cytonuclear, mitochondrial, and ER-targeted proteins

To study NAĆs association with translating ribosomes, we performed metagene analyses on the SeRP_ribo_ and SeRP_NC_ datasets. Given NAC’s roles in ER translocation and mitochondrial import^3,15,16,38–40^, nascent proteins were categorized by cellular location: (i) cytonuclear proteins (nucleus, cytoplasm), (ii) ER-targeted proteins (ER lumen, ER membrane (ERM), plasma membrane (PM)), and (iii) mitochondrial proteins (matrix, intermembrane space, membrane).

In the SeRP_ribo_ dataset, metagene profiles for cytonuclear and mitochondrial proteins showed uniform NAC enrichment across CDS (**Figures 2A and 2B**), confirming NAĆs NC-independent binding and in agreement with the individual examples described above (**Figures 1B, 1C S1G-S1I**). Conversely, the SeRP_NC_ dataset revealed an NC-dependent binding pattern (**Figures 2A and 2B**) with minimal interactions early in translation, followed by a steady increase that plateaus after the synthesis of approximately 70 residues. Over 70% of nascent cytonuclear and mitochondrial proteins had at least one high-confidence NAC binding period, classifying them as NAC interactors (**Table S1**). These data confirm that NAC ubiquitously associates with ribosomes translating cytonuclear and mitochondrial proteins.

**Figure 2:**
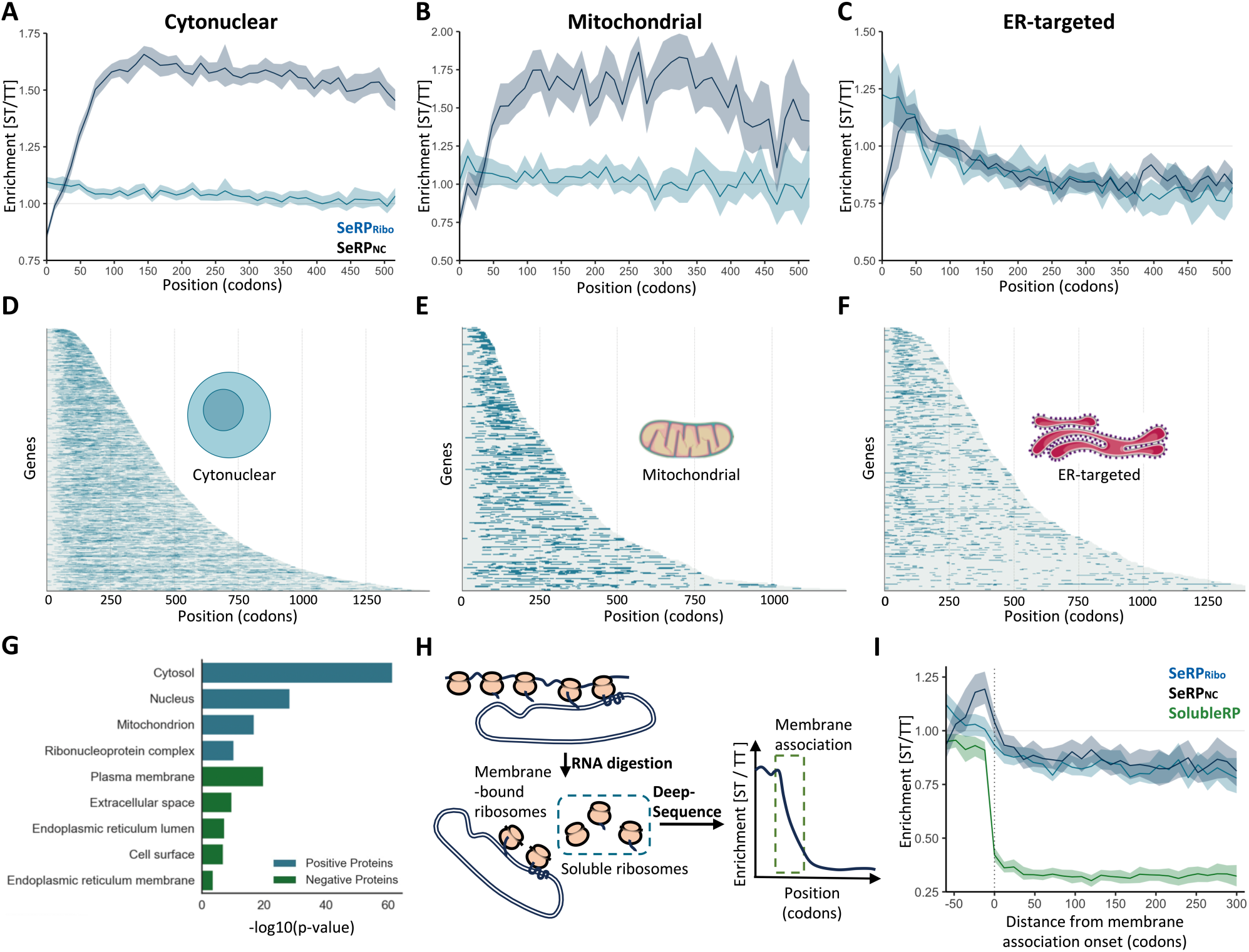
NAC binding patterns in nascent cytonuclear, mitochondrial and ER targeted proteins. (**A-C**) Metagene profiles showing NC-dependent (SeRP_NC_) and NC-independent (SeRP_Ribo_) ST/TT enrichment along coding sequences of cytonuclear, mitochondrial and ER-targeted proteins. Metagenes include 6824, 536 and 1393 genes with coverages > 0.1 reads/codon, respectively. Shadowed areas indicate 95% confidence intervals. (**D-F**) Heat maps of NC-dependent NAC binding periods (blue) in cytonuclear (D), mitochondrial (E) and ER translocated (F) proteins. Heatmaps include 3275, 316, 515 genes, respectively. (**G**) Gene ontology analysis by cellular component of positive (at least one NAC binding period) and negative proteins. (**H**) Schematic representation of the SolubleRP pipeline used to detect co-translational membrane association. (**I**) Metagene profile showing NC-dependent, NC-independent and SolubleRP enrichments along coding sequences of high confidence co-translationally translocated proteins aligned to the onset of membrane association. Shadowed areas indicate 95% confidence intervals.

In contrast, ER-targeted nascent proteins exhibited a length-dependent depletion in the SeRP_ribo_ dataset, indicating that NAC dissociates as translation progresses (**Figure 2C**). In the SeRP_NC_ dataset, NAC binding increased as NCs emerge from the tunnel exit, peaking around residue 50 before decreasing. A divergence for ER-targeted proteins was also evident in the distribution and prevalence of NAC binding periods, being less frequent and biased towards shorter NCs compared to cytonuclear and mitochondrial nascent proteins (**Figures 2D, 2E and 2F**). The duration of NAC binding periods was also longer for cytonuclear and mitochondrial proteins (median width: 26 codons, **Figure S3A**) compared to ER-targeted proteins (median width: 20 codons). Only 43% of nascent ER-targeted proteins were classified as NAC interactors (**Table S1**). This localization dependency was confirmed by Gene ontology analysis (**Figure 2G**): non-interactors were enriched in ER-targeted proteins, while interactors were enriched in cytonuclear and mitochondrial categories.

The NAC interactome for ER-targeted proteins suggests that NAC initially associates with NCs but is later displaced during co-translational ER translocation (**Figure 2C**). We confirmed this using an adaptation of ribosome profiling, termed SolubleRP, which measures ribosome depletion from the soluble fraction of a cellular lysate (after nuclease digestion of polysomes) as a proxy for ER translocation (**Figure 2H**). This approach provided detailed insights into ribosome engagement with membranes, revealing a NC-dependent depletion of soluble monosomes translating ER-targeted proteins (**Figure S2A**). We identified the onset of membrane targeting for a set of high-confidence co-translationally ER-targeted substrates (**Table S2**) and found a strong correlation with the emergence of signal peptides (SPs) and transmembrane domains (TMDs) (**Figure S2B**). Aligning the SeRP_Ribo_ and SeRP_NC_ datasets to the onset of membrane association revealed that NAC binds nascent chains prior to membrane targeting, but rapidly dissociates thereafter (**Figure 2I**). This *in vivo* observation aligns with previous reports of NAC’s participation in SRP-mediated targeting, based on stalled ribosome-NC complexes *in vitro*^3^. The metagene analysis was supported by individual examples, including the secreted protein B2M and multi-pass membrane proteins TSPAN3 and DEGS1 (**Figure S2C**). To further evaluate NAC’s involvement in ER targeting, we examined its binding relative to SP and TMD emergence. We found that exposure of 25% and 45% of co-translationally targeted SPs and TMDs coincided with NAC binding periods (**Figures S3B and S3C**). However, no statistically significant differences in the overall physicochemical properties of SPs and TMDs were observed between NAC-bound and NAC-unbound groups (**Figures S3D and S3E**). Thus, while NAC binding is prevalent among co-translationally targeted SPs and TMDs, it was not universal, suggesting the presence of alternative, NAC-independent targeting routes.

In addition to its role in ER targeting, NAC has also been proposed to mediate NC recruitment to the mitochondrial surface^16,38–40^. Unlike ER-targeted proteins (**Figures 2C and 2F**), NAC binding to nascent mitochondrial proteins showed only a modest decrease during translation (**Figures 2B and 2E**), in agreement with mitochondrial precursors being primarily synthesized in the cytosol before translocation^41^. To investigate NAC’s involvement in mitochondrial import, we analyzed its binding to cleavable mitochondrial targeting sequences (MTS), which direct the import of inner mitochondrial membrane (IMM), intermembrane space, and matrix proteins. 30% of the analyzed MTS were bound by NAC upon their emergence from the ribosomal tunnel exit (**Figure S4A**). Notably, proteins with NAC-bound MTS were enriched in IMM proteins (p-value < 0.05, CC-GO enrichment), with 45% of IMM-targeted MTS and 24% of matrix-targeted MTS showing NAC association (**Table S3**). Driven by this observation, we analyzed NAC binding to TMD-containing mitochondrial proteins in more detail, including IMM and outer mitochondrial membrane (OMM) proteins. NAC binding periods coincided not only with MTS emergence, but also with TMD emergence from the exit tunnel in both OMM and IMM proteins (**Figure S4B; Table S4**), predominantly in polytopic membrane proteins (**Figure S4D**). Unlike ER-translocated proteins, TMD binding did not exhibit an N-terminal bias (**Figure S3C**) and showed no significant differences in the physicochemical properties of bound versus unbound TMDs (**Figure S4C**). In support of the functional relevance of these interactions, the polytopic OMM proteins TSPO and FUNDC1 (enrichment profiles in **Figures S4E and S4F**), whose mitochondrial import is impaired upon NAC depletion^42^, were included in the NAC-bound group (black arrows). In contrast, the tail-anchored proteins CISD1 and MAVS (red arrows), which are unaffected by NAC depletion, were not bound by NAC. The TSPO enrichment profile revealed a single, specific NAC-binding event at the second emerging TMD (**Figure S4E**), suggesting that NAC’s role in TSPO import is limited to a short-lived interaction with a defined transmembrane region. Overall, our data support a potential role for NAC in the biogenesis of both IMMs and OMMs, extending its previously reported involvement in OMM biogenesis^42^.

In summary, our profiling analysis provides a proteome-wide view on NAC’s roles in ER targeting and mitochondrial import, uncovering its *in vivo* substrate pools and major binding principles. However, we also observed that NAC’s association with mitochondrial proteins extended beyond N-terminal MTS- and TMD-containing substrates (**Figure 2E; Tables S1 and S4**). These NAC binding periods within nascent mitochondrial precursor proteins outside emerging MTS and TMD sequences shared features similar to those found in cytonuclear proteins (see below). This suggests multiple roles for NAC in mitochondrial protein import, potentially including co-translational chaperoning.

### NAC recognizes emerging domains of cytonuclear proteins

We next investigated the functional significance of NAC binding to nascent cytonuclear proteins. Given that NAC binding often coincided with the emergence of annotated protein domains in individual examples (**Figures 1B, 1C and S1G-S1I**), we examined this correlation at a proteome-wide scale. Across the proteome, emerging cytonuclear protein domains indeed showed increased NAC interactions compared to flanking regions (**Figure 3A**), a pattern also observed in mitochondrial proteins (**Figure 3B**) but not in ER-targeted proteins (**Figure S5A**). Metagene NAC binding profiles showed a sharp initiation at the beginning of domain exposure, followed by a gradual decline as the domain is further translated. Notably, NCs with early-emerging domains exhibited earlier NAC binding (**Figure 3C**), emphasizing the importance of the strong correlation between domain emergence and NAC engagement. In agreement, analysis of NC features at the tunnel exit during NAC binding periods revealed a correlation with protein domain characteristics, including a high hydrophobic and aromatic residue content, increased α-helical and β-sheet structures, reduced predicted disorder, and lower accessible surface area (ASA). This suggests that NAC binds elements destined to be buried in the final structure (**Figure S5B**). Long disordered regions had fewer NAC binding peaks (**Figures 1B, 1C and S1G-S1I**), a trend also seen in non-globular and coiled-coil proteins lacking folded domains (**Figures S5C and S5D**).

**Figure 3:**
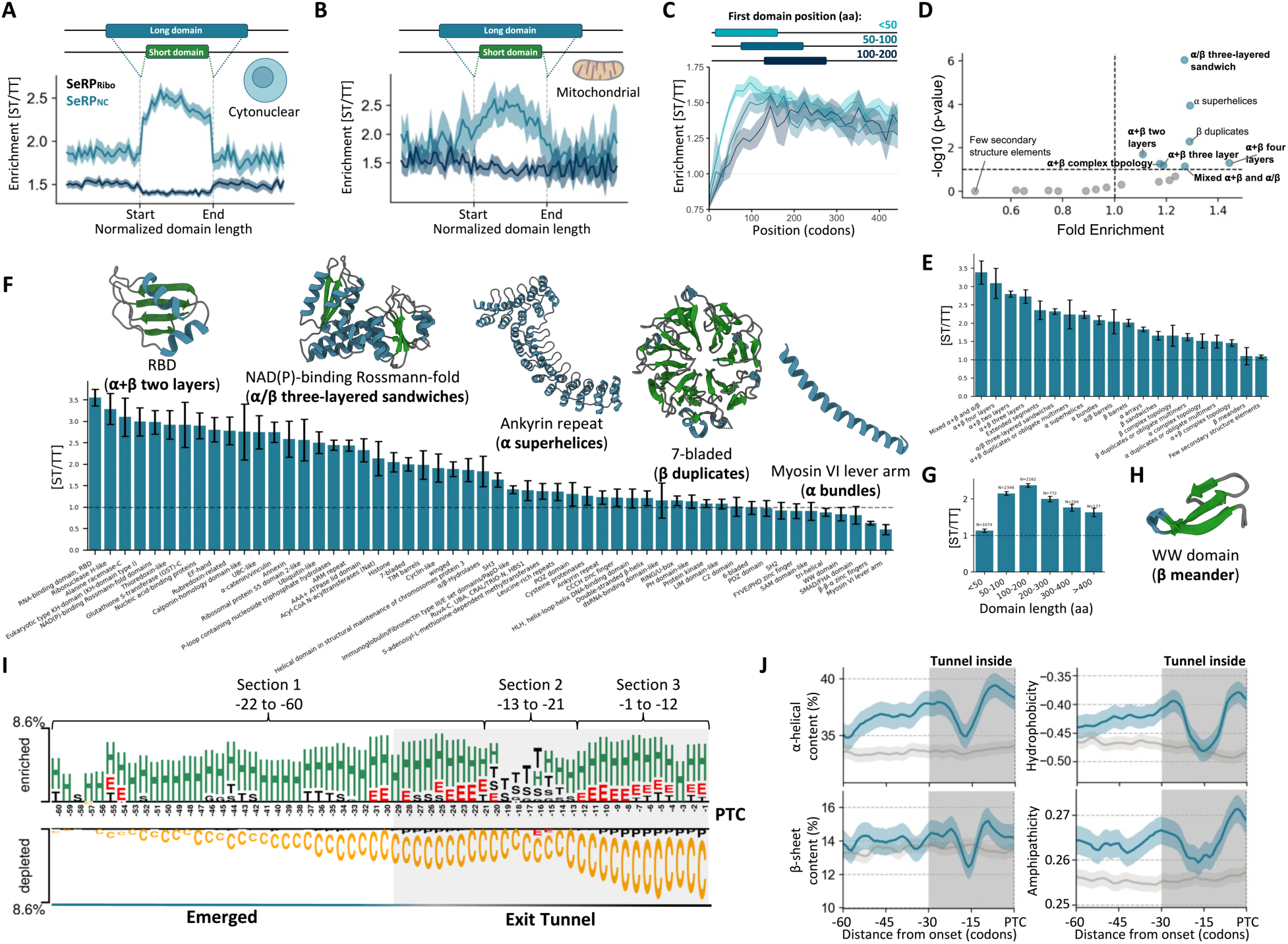
Molecular determinants of NAC binding to nascent cytonuclear and mitochondrial proteins. (**A-B**) Length-normalized ST over TT enrichment of cytonuclear (A) and mitochondrial (B) proteins within domains and in flanking regions for SeRP_Ribo_ (black) and SeRP_NC_ (blue) interactions. (**C**) Metagene profiles of proteins categorized by the position of their most N-terminal domain. Domain positions are segmented into three groups: proteins with the first domain starting at positions <50, 50–100, or 100–200. Metagenes include 3250, 1115 and 805 proteins, respectively. Shadowed areas indicate 95% confidence intervals. (**D**) Frequency enrichment analysis of domains bound by NAC (i.e: containing at least one binding period). (**E**) Total ST/TT enrichment on different ECOD domain architectures. (**F**) Total ST/TT enrichment on different ECOD domain topologies. Underrepresented topologies (<20 examples in our dataset) were excluded. Structural examples of different topologies are shown (architecture in brackets) (PDBs: 8csv, 1hd0, 1a4i, 1oe9 and 1i2m). (**G**) Total ST/TT enrichment of protein domains according to their length. (**H**) Example of a β meander domain (PDB: 1ywj). (**I**) Position-specific enrichment of the DSSP annotated secondary structure in ribosome-proximal 60 residues at the onset of NAC binding in cytonuclear proteins. Annotation following DSSP classes; H: helix, E: strand, T: turn, S: bend, C: Coil, G:3_10_-helix, P: pi-helix. (**J**) Metagene profile of helical content (DSSP annotated from Alpha-fold structures), β-sheet content (DSSP annotated from Alpha-fold structures), hydrophobicity (Kyte-Doolittle)^64^ and amphipathicity measured as the hydrophobic moment of the helix^65^ aligned to the onset of NAC binding periods in cytonuclear proteins. Gray line corresponds to a randomized background control. Shadowed areas indicate 95% confidence intervals.

NAC binding profiles varied strongly between domains, for instance showing differences in the number of binding peaks within a single domain (**Figures 1B, 1C and S1G-S1I**), prompting an analysis of specific domain architectures linked to NAC binding (**Figure 3D**). NAC-bound domains were enriched in mixed α and β architectures, α superhelices, and β duplicates compared to other domain architectures. Local NAC enrichment within domains mirrored this trend, with strong NAC binding to mixed α and β architectures (**Figure 3E**). α superhelices were the main α architecture bound by NAC, while β duplicates show lower enrichment values. Domains with minimal secondary structure elements were notably depleted in NAC binding (**Figures 3D and 3E**). Examining NAC preferences for domain topologies revealed a similar trend (**Figure 3F**), with domains like RNA-binding domains (RBD; α+β two-layered architecture) and NAD(P)-binding Rossmann-fold (α/β three-layered sandwiches) showing strong NAC binding, while Myosin VI lever arm (α bundles) and WW domains (β meander) showing reduced binding.

Overall, these findings suggest a role of NAC in the biogenesis of folded domains, congruent with NAC being a co-translationally acting chaperone for cytonuclear proteins. Each domain architecture has a distinct NAC binding propensity, suggesting an evolutionarily selected chaperone activity. The stronger binding of NAC to mixed α and β architectures likely relates to their complex topology and folding needs, while small domains under 50 residues with lower folding complexity showed less NAC binding (**Figure 3G**), exemplified by lower binding in SH2 (α+β two layers), tri-helical (α arrays), β-β-α zinc fingers (few secondary structure elements) and WW domains (β-meander; **Figure 3H**).

### Molecular determinants of NAC binding to cytonuclear proteins

We analyzed the physicochemical features of nascent cytonuclear proteins at the onset of NAC binding. Given the small size of NAĆs β-barrel domain, its position at the tunnel exit, broad substrate range and ability to bind protein segments within emerging domains (**Figure 3A**), we surmised that NAC recognizes primary sequence or secondary structure motifs close to the tunnel exit, rather than complex tertiary folds. Supporting this, NAC binding spans a median of 26 codons (**Figure S3A**), shorter than full protein domains (**Figures 1B, 1C and S1G-S1I**). To investigate secondary structure preferences, we examined the position-specific enrichment of AlphaFold-based DSSP secondary structure annotations in the 60 most recently translated residues at the start of NAC binding (**Figure 3I**). Both α and β secondary structures were enriched, while disorder decreased compared to a randomized control, consistent with NAC binding to folded domains. NAC binding also correlated with the translation of secondary structure elements, both α-helices and β-sheets, with an intermediate section enriched in connecting loops (**Figure 3I**, sections 2 and 3). While these motifs are buried in the ribosomal exit tunnel at the onset of NAC binding, they become partially or completely exposed during the binding period, suggesting that NAC binding is maintained during the emergence of such secondary structure motifs.

Focusing on the NC segments accessible to NAC at binding onset, we found a clear correlation between the start of NAC binding and the emergence of α-helices (**Figure 3I**). Exploring the physicochemical properties of these segments at the metagene level revealed that NAC binding coincides with the emergence of regions enriched in hydrophobicity, α-helical propensity and amphipathic characteristics (**Figures 3J and S6A**). β-sheet enrichment was lower, indicating that hydrophobicity and high helical propensity are the primary binding criterium, consistent with the preference for α superhelices over β duplicates. A similar position-specific analysis of primary sequences did not yield a sequence-specific binding motif (**Figure S6B**), but amino acid compositions aligned with the observed secondary structure propensities; residues disrupting secondary structures (G, P) were less prevalent in sections 1 and 3, while aliphatic and aromatic (ILVMFY) residues were enriched. Conversely, G and P were common in section 2, aiding rigid loop formation.

These results indicate that NAC interactions are guided by the conformational properties of NCs rather than a specific primary sequence, supporting its broad substrate binding. The enrichment of secondary structure elements connected by short loops suggests that NAC preferentially targets motifs destined to form well-folded, compact local structures. This explains NAĆs reduced binding to extended structures like the long helix in Myosin VI level arm domains or extended coil-coil regions (**Figures 3F and S5D**), while also accounting for its preference for α superhelices, characterized by densely packed α-helices connected by loops. Overall, these data hint that NAC participates in the biogenesis of minimal tertiary folds composed of (α/β)-loop-(α/β) motifs.

### Direct observation of dynamic NAC interactions with the elongating nascent chain

SeRP provides a proteome-wide view of NAC-NC interactions *in vivo*, but the cellular complexity limits detailed mechanistic insights. To explore the molecular basis of NAC–NC interactions, we used a reductionistic *in vitro* approach with smFRET measurements based on Total Internal Reflection Fluorescence Microscopy (TIRFM) (**Figure S7A**). RPL4 was selected as a model substrate (**Figures 4A and S7B**), as it is a highly translated protein suitable for generating stalled ribosome-NC complexes (RNCs) and NAC-bound ribosome structures translating RPL4 are available^4^. The RPL4 globular domain adopts an α/β three-layered sandwich architecture (**Figure 4A**), with two segments separated by an intradomain linker. Our SeRP data indicate NAC binds both RPL4 segments but not the linker (**Figure 1C**).

**Figure 4:**
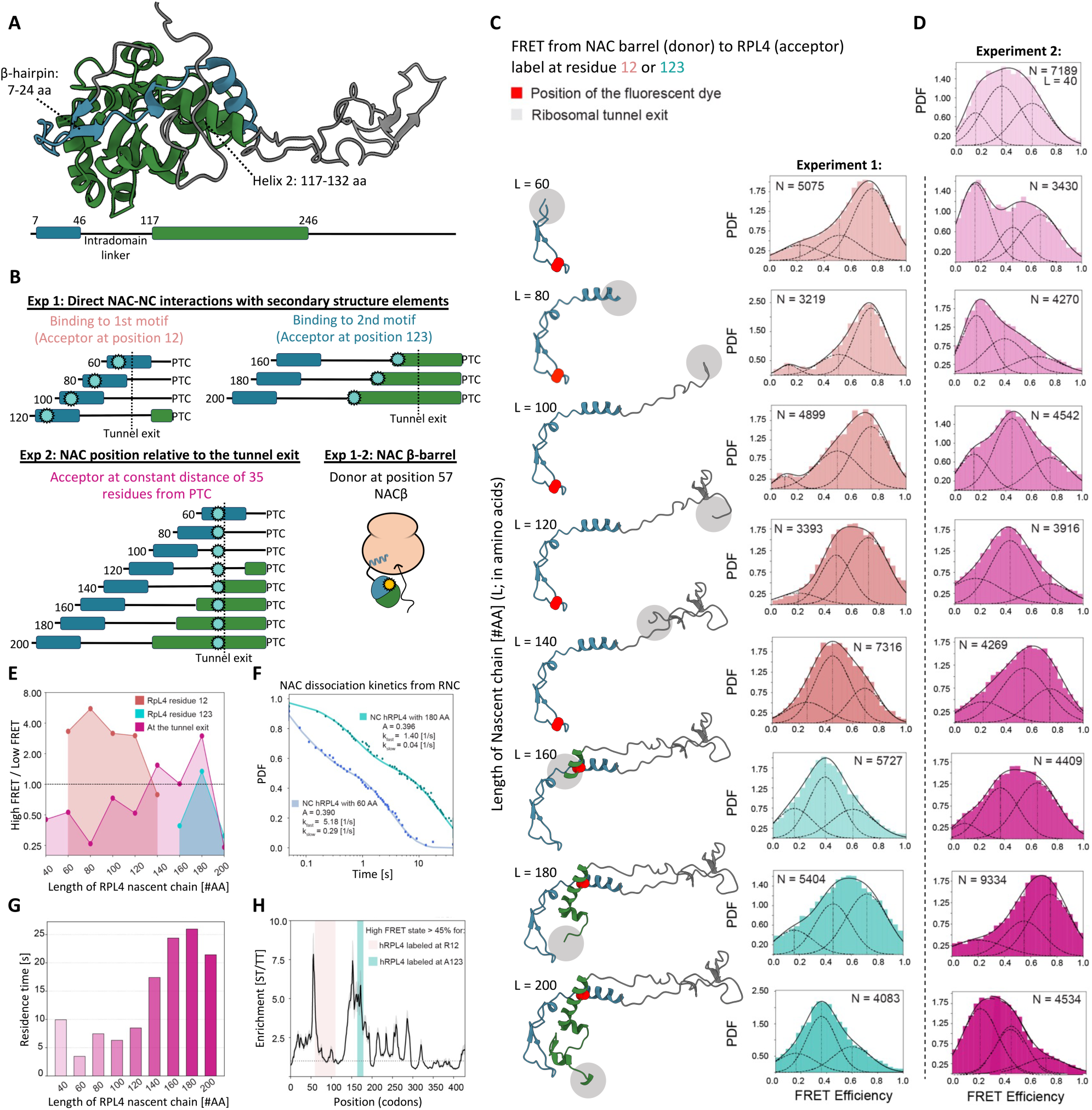
Single molecule FRET analysis of NAC interaction with ribosomes and RPL4 nascent chains. (**A**) AlphaFold model showing RPL4 globular domain organization. Blue and green colors indicate the N- and C-terminal segments of the RPL4 globular domain, respectively. **B**) Schematic representation of the lengths of the RpL4 NC and the position of the fluorophore (cyan circles) in the RPL4 RNCs analyzed. (**C**) smFRET histograms for energy transfer between a donor dye labeled in the β-barrel domain of NAC and an acceptor dye labeled at residue 12 or 123 in the RPL4 NC. The lengths of the RPL4 NCs are indicated. The AlphaFold model of the region of RPL4 outside the exit tunnel at each NC length is depicted on the left, with dye-labeled residues 12 or 123 highlighted in red. The *grey* sphere indicates the position of the ribosome tunnel exit. (**D**) smFRET histograms for energy transfer between a donor dye in the β-barrel domain of NAC and an acceptor dye positioned at the ribosome exit site. In (C) and (D), ‘N’ denotes the number of measurements used to construct the histogram. The data were fit to the sum (solid lines) of three Gaussian functions (dashed lines). (**E**) Summary of the distribution of NAC in high FRET populations at different NC lengths for the FRET pair with the acceptor dye at RPL4 residue 12 (salmon), RPL4 residue 123 (teal), and at the ribosome exit tunnel (pink). The number of molecules with FRET > 0.6 are summed and divided over those with FRET < 0.4. (**F**) Representative kinetic traces showing the dissociation of NAC from RNCs with NC lengths of 60 and 180 residues. The data were fit to the sum of two exponential functions, which gave the dissociation rate constants for the fast- and slow-dissociating populations (*k*_fast_ and *k*_slow_) and the fraction of the fast-dissociating population (A). (**G**) Summary of the residence times of the slowly dissociating NAC fractions on the RPL4 RNC at different NC lengths. (**H**) NC lengths at which NAC-NC interactions were detected in smFRET data are mapped onto the NAC profiling data on RPL4.

To directly observe the ribosome and NC interactions of NAC and to understand whether they compete or cooperate with one another, we labeled RNCs with the acceptor dye Atto647N at various positions in the NC (**Figure 4B**) and immobilized them on imaging slides via a biotin moiety at the 5’ end of the mRNA (**Figure 7A**). NAC was labeled with the donor dye Cy3B at residue 57 of the β-barrel domain. To probe the interaction of NAC with the two segments in the RPL4 globular domain (Experiment 1), Atto647N was incorporated at residue 12 or 123 of RPL4 (**Figures 4A, 4B and 4C**). To probe the ribosome interaction of the NAC barrel domain near the exit tunnel (Experiment 2), Atto647N was incorporated at a defined distance of 35 residues from the NC C-terminus to ensure that it was positioned at the tunnel exit across all the RNCs with varying NC lengths (**Figures 4B and 4D**). NAC binding to immobilized RNCs was assessed via fluorescence colocalization and FRET between the dye pair under TIRFM (**Figure S7A**). The TIRFM setup also allows direct measurement of NAC interaction lifetimes, by moninoring the duration of NAC association (colocalization) with surface-immobilized RNCs (**Figure S7A**).

Hidden Markov Modeling (HMM) of smFRET traces revealed at least three FRET states, with low, medium, and high FRET efficiencies, for both experimental probe configurations (**Figures S8A and S8B**). At a NC length of 40 residues, the smFRET histogram for NAC with the acceptor dye at the tunnel exit (Experiment 2) displayed a broad distribution, with ∼50% in medium FRET and 35% in high FRET states (**Figure 4D**, top panel). The high FRET state is assigned to the conformation in which the NAC β-barrel domain docks near the exit tunnel^3^, while the medium/low FRET states are attributed to conformations where the NAC barrel dissociates from its ribosome docking site, either remaining free or engaging in alternative interactions. As only the N-terminal ∼5 residues of RPL4 have emerged from the exit tunnel at this NC length, this histogram serves as a ‘baseline’ for NAC barrel distribution at the exit tunnel (Experiment 2) in the absence of significant NC interactions.

As the RPL4 NC elongates, changes in smFRET distributions indicate two interaction stages between the NAC barrel and the NC. The first occurs at NC lengths of 60-80 residues, where the NAC barrel domain primarily occupies a low FRET state with the acceptor dye at the tunnel exit (Experiment 2) (**Figures 4D, 4E, and S8D**), suggesting that it moved away from the ribosome due to competing interactions with the NC. Consistently, the high FRET state dominated the smFRET histograms with the acceptor dye at RPL4 residue 12 at these NC lengths (Experiment 1) (**Figures 4C, 4E, and S8D**). The anti-correlation of these two FRET pairs (**Figure 4E**) suggests a preferential interaction of the NAC β-barrel with the N-terminal segment of the RPL4 globular domain, which outcompetes ribosome binding. As the NC elongates, the high FRET population with the probe at RPL4 residue 12 (Experiment 1) decreased until it became a minor state at 140 residues (**Figures 4C, 4E and S8C**), indicating the dissolution of NAC interaction with this NC region.

The second stage of NC interaction occurred when the RPL4 NC exceeded 160 residues, marked by an increased kinetic stability of the NAC-RNC interaction (**Figures 4F and 4G**). As reported previously^42^, NAC displayed biphasic dissociation kinetics from the RNC, with rate constants of 0.29 s^-1^ and 5.2 s^-1^ for the slow- and fast-dissociating populations, respectively, at a NC length of 60 residues (**Figure 4F**). While the distribution of these populations and the stability of the fast-dissociating population changed modestly across the NC lengths (**Figure S9**), the lifetime of the slow-dissociating population increased from 4-10 sec at shorter lengths to ∼25 sec at lengths exceeding 160 residues (**Figures 4F and 4G**), indicating enhanced NAC-RNC interactions. At a NC length of 180 residues, high FRET states were prominent in both smFRET histograms of NAC paired with the acceptor dye at the ribosome tunnel exit and at residue 123 in the C-terminal segment of the RPL4 globular domain (**Figures 4C- 4E**). At this stage, favorable NAC interactions with both the surface of the ribosome and the nascent chain of RPL4 likely contribute to the enhanced NAC-RNC stability.

Collectively, the smFRET data show that the NAC β-barrel domain dynamically samples the evolving NC during translation, establishing interactions with defined RPL4 regions at specific stages of translation. These interactions can modulate the position of the NAC β-barrel at the ribosomal exit tunnel, either displacing it or bringing it into closer proximity. The two detected NAC interaction regions on the RPL4 NC were in agreement with the SeRP_NC_ data (**Figure 4H**). As the smFRET studies were carried out with purified complexes in the absence of other cellular factors, this agreement suggests that the observed preference of NAC for NC features does not rely on other factors. NAĆs longer interactions with the N-terminal region of RPL4 during translation in smFRET compared to SeRP may be attributed to additional factors that compete for interaction *in vivo*, such as MetAP1, NatA or other chaperones, leading to earlier NAC release from the N-terminal region during translation *in vivo*. In agreement with this possibility, the intradomain loop of RPL4 is known to be co-translationally bound by an RPL4-specific chaperone in yeast^43^.

Both NAC-interacting regions of RPL4 correspond to local structural elements within its globular domain that has yet to complete translation and folding (**Figure 4A**). The initial NAC interaction occurred with the fully emerged N-terminal β-hairpin in the N-terminal segment of RPL4, which will also contact a helix-turn-helix region of the globular domain that has not yet emerged at this stage of translation. The second interaction occurred when helix 2 at the core of the partially translated globular domain of RPL4 fully emerged from the ribosome (**Figure 4C**). These observations are in agreement with the NAC interaction patterns observed in SeRP and support the notion that NAC recognizes secondary structural elements or minimal tertiary folds that form locally at this early stage of folding, pointing to an early chaperone function of NAC.

### Structural insights into NAC interaction with nascent chains

To understand the molecular basis of NAC binding to NCs, we determined the structure of NAC in complex with RPL4 RNC using cryo-EM (**Figure 5A**). While previous studies determined NAC-RNC structures in complex with different ribosome-associated factors^2–4,13,44^, direct interactions between NAC and NCs have not been observed. Driven by our observation in SeRP and smFRET studies, we generated stalled RNCs with NCs comprising the first 158 amino acids of RPL4 followed by the Xbp1u translational arrest peptide^45^, which mimics an 180 amino acid-long RPL4 NC. At this length, high FRET signals were detected between the NAC β-barrel and both the acceptor dye at position 123 of the NC and the dye at the tunnel exit, and RNC-NAC binding peaked in kinetic stability.

**Figure 5:**
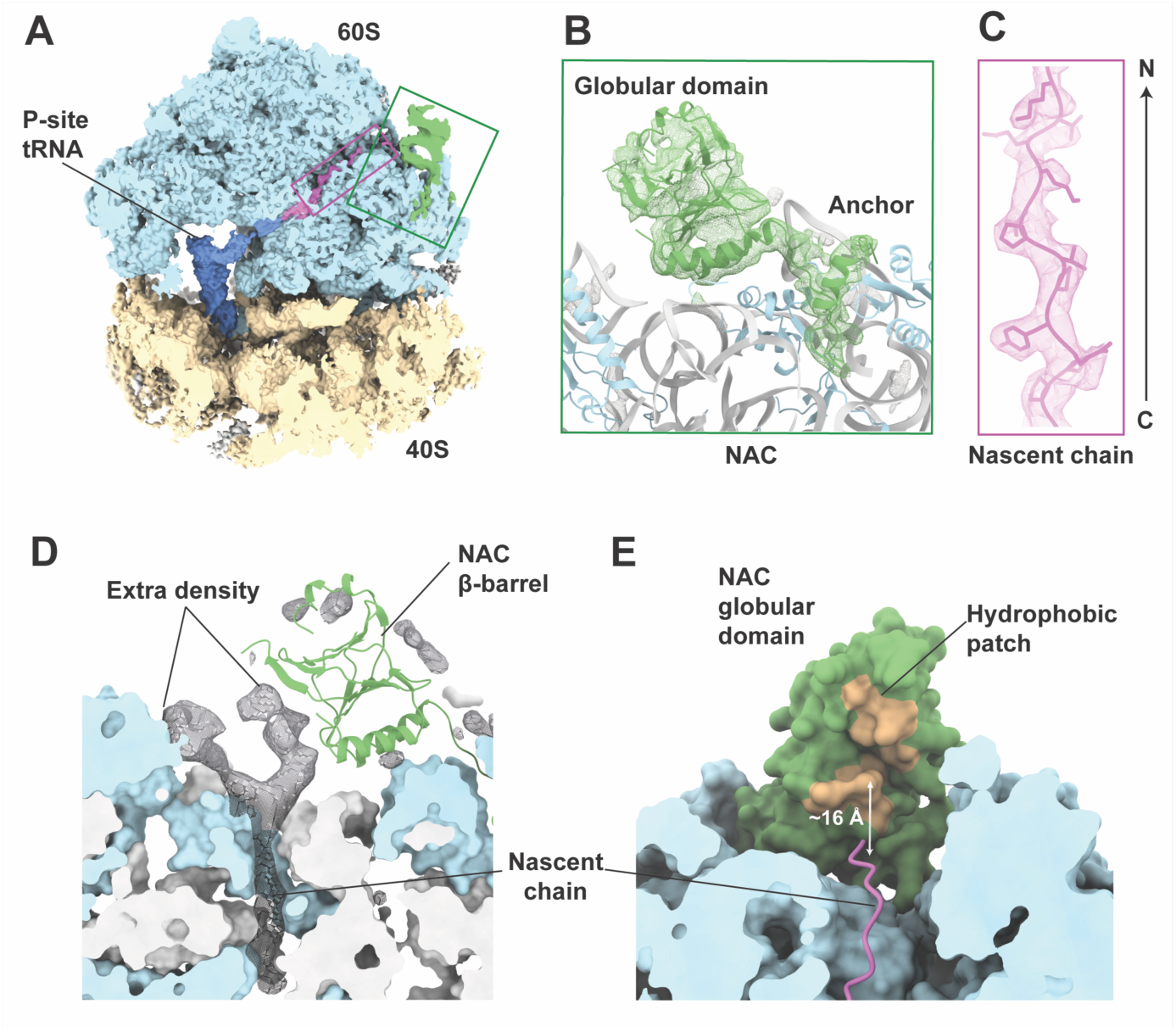
Cryo-EM analysis of RPL4 ribosome-nascent chain (RNC) complexes with NAC. (**A**) Cross-section overview of the cryo-EM map of RPL4 RNC in complex with NAC, low-pass filtered to 4 Å. The 60S proteins, 28S, 5S and 5.8S rRNAs are shown in light blue, 40S proteins and the 18S rRNA in yellow, the NAC heterodimer in green, the P-site tRNA in dark blue, and the nascent chain in pink. Rectangles indicate the position of close-up views in B-C. (**B-C**): Close-up views of NAC (B) and the nascent chain in the polypeptide exit tunnel (C). The difference density between the cryo-EM map and the 80S model, low-pass filtered to 4 Å, is shown in green (B), while the 2.8 Å difference map is represented as pink mesh (C). 60S proteins are shown in light blue, rRNA in gray, NAC in green and the nascent chain in pink. (**D**) Continuation of the nascent chain density outside the tunnel exit. The difference density between the cryo-EM map and the 80S/NAC model, low-pass filtered to 6 Å, is shown as black mesh. 60S proteins are shown in light blue, rRNA in light gray, and NAC in green. (**E**) Bottom view of the NAC β-barrel in the RPL4 RNC/NAC complex, highlighting the hydrophobic patch (orange) on NAC facing the tunnel exit. The approximate distance from the tunnel exit to the hydrophobic patch is indicated. 60S proteins and rRNA are shown in light blue, NAC in green/orange, and the nascent chain in pink.

The cryo-EM map shows NAC bound at the polypeptide tunnel exit of RPL4 RNCs at the same position as described previously (**Figure 5B**)^3,4^, with the N-terminal basic ribosome anchor of NACβ binding to eL19 and eL22, and two positively charged α-helices of the NAC β-barrel domain contacting rRNA. The nascent polypeptide chain could be traced throughout the exit tunnel (**Figure 5C**). Additionally, we observed a continuation of the nascent chain density outside the tunnel exit at lower thresholds (**Figure 5D**). The extra density was observed along the 28S rRNA lining the polypeptide exit tunnel and contacted 5.8S rRNA on the surface of the ribosome, as well as the entrance to the NAC β-barrel. This surface of NAC presents a patch of hydrophobic residues, which are contributed by both NACα and NACβ, at short distance from the tunnel exit (**Figures 5E and S10A**). The location of the nascent chain density outside the tunnel exit was reproducible in two independently obtained cryo-EM maps (**Figure S10B**). Consistent with a stabilizing interaction between NAC and the NC, the classification for the conformation of the NAC globular domain of the cryo-EM map revealed a more defined NC density outside the exit tunnel (**Figure 5D)**, compared to the map prior to classification (**Figure S10B**). Additionally, the NAC β-barrel domain was resolved to a higher resolution (∼5 Å) than in previous studies^3,4^, which may also be rationalized by a more uniform positioning of the NAC β-barrel domain due to stabilizing interactions with the NC.

The accumulation of a NC density next to the NAC globular domain is consistent with the high FRET state (**Figure 4D**) and SeRP enrichment (**Figure 1C**) at this NC length, with multiple NAC lysines present within DSP spacer length (12 Å) of the NC (**Figure S10C**). Residue 57 of NACβ, where the FRET donor was placed, was also in close proximity to the NAC interaction surface. Notably, the NC extension outside the tunnel exit was absent in previous studies using shorter RPL4 chains^4^ and other NCs^2,3,13,44^, indicating NC-specific interactions with the NAC globular domain, likely mediated by the hydrophobic patch on the NAC β-barrel.

The smFRET data (**Figure 4**) revealed a close proximity between residue 123 of RPL4 and the NAC binding surface, suggesting that the observed density may originate from NC regions extending from the last assigned residue toward the N-terminus (residues 123–151). Thus, while the exact conformation of the NC interacting with NAC remains unresolved, the cryo-EM density and smFRET data together suggest that the NC adopts a partially collapsed state. This is consistent with NAC’s preference for loop-containing motifs, which can adopt such a compacted structure, as observed in the SeRP_NC_ data. However, the proximity of the NAC β-barrel domain to the tunnel exit would likely prevent higher-order folding. NAC may therefore provide an interaction surface that facilitates early formation of an initial folding nuclei as the NC emerges from the polypeptide tunnel exit.

### NAC binding promotes protein folding

We next aimed to investigate NAĆs possible role in protein folding. While the study of nascent chain folding dynamics at the human ribosome is challenging, it is feasible to probe how NAC interactions affect the folding of a polypeptide chain. For this purpose we used single molecule optical tweezers to investigate Maltose Binding Protein (MBP) as a model substrate^46^. MBP is well-characterized for studying chaperone guided protein folding by tweezer experiments, including trigger factor (TF), DnaK/Hsp70, and GroEL^47–49^ thus providing a robust benchmark for the mechanistic dissection of NAC’s role in the folding of proteins with mixed α and β architecture.

We assessed MBP folding efficiency in the absence or presence of NAC. Single MBP molecules were tethered between two laser-trapped polystyrene beads using DNA handles (**Figure 6A**, see Methods). After mechanically unfolding the tethered protein by moving one bead away from the other, we subjected MBP to repeated cycles of relaxation (by bringing the beads together), waiting for 5 s in the absence of applied force to allow refolding, and stretching (by separating the beads again) (**Figures 6A and 6B**). The polypepetide chain either folded, as evidenced by characteristic unfolding of the MBP core during subsequent stretching, or remained unfolded, as shown by a lack of unfolding features during stretching (**Figure 6B**, right), as previously demonstrated^50^. A few C-terminal alpha-helices could dock onto the core structure, and conversely unfold in a stepwise or gradual transition at about 10 pN during stretching, such that the core state remains (**Figure 6B**, left). Because formation of the MBP core state is the rate-limiting step in folding, we quantified the fraction of cycles showing MBP core formation. Remarkably, the presence of 5 µM purified NAC heterodimers in solution significantly enhanced this MBP core refolding frequency from 0.51 in the absence of NAC to 0.90 with NAC present (**Figure 6C**). Given that protein folding involves the probabilistic crossing of an energy barrier, these observations indicate that NAC effectively lowers this folding energy barrier and hence promotes the folding process^49,51^.

**Figure 6:**
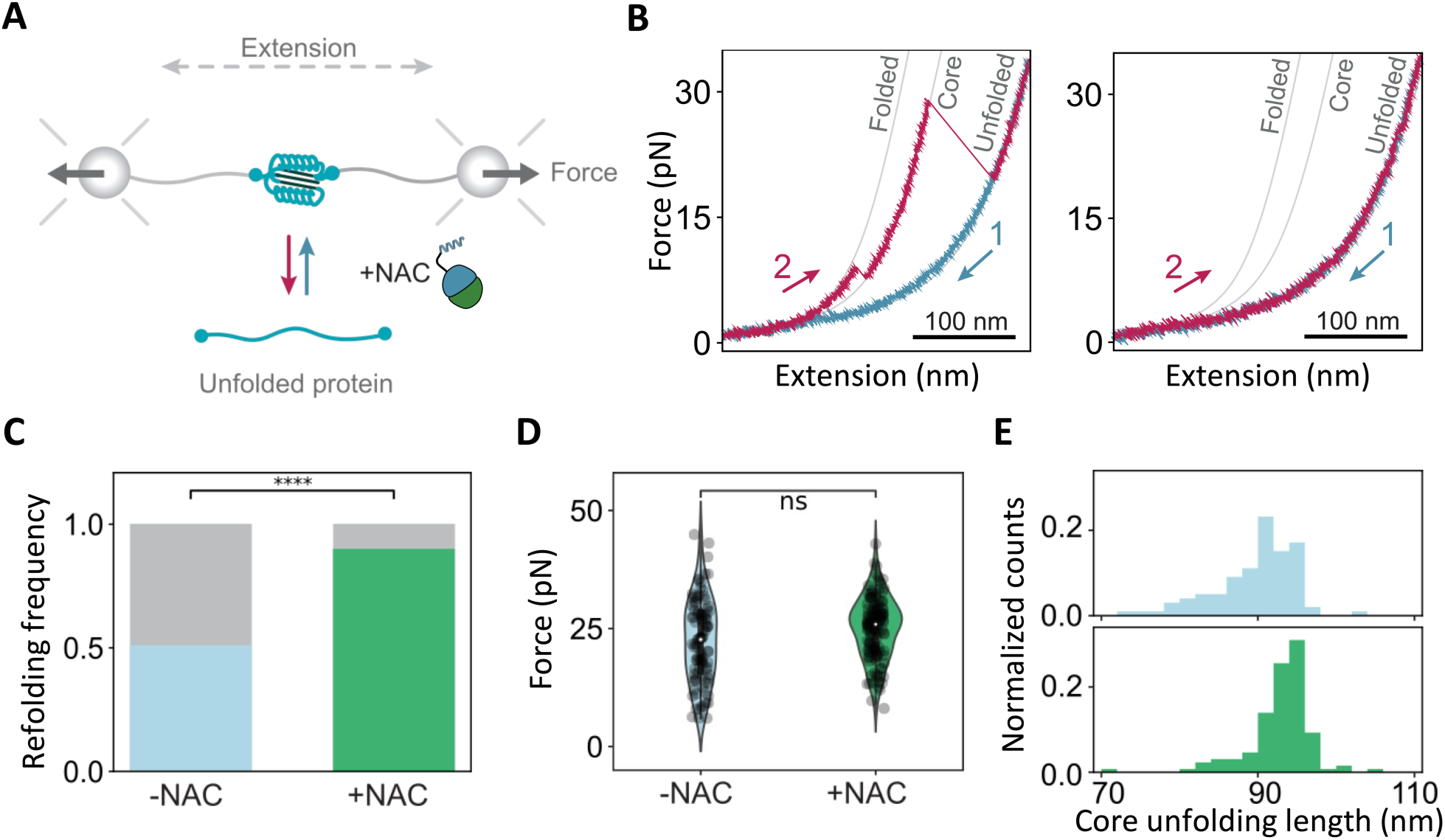
NAC facilitates native folding of a model protein. **(A)** Schematic illustration of the optical tweezers assay. **(B)** Representative force-extension curves of folded (left panel) and unfolded MBP (right panel), recorded in the absence of NAC. Following the relaxation of unfolded MBP molecule (1) and waiting at 0 pN for 5 s, the stretching curve (2) displays a step corresponding to unfolding of the MBP core structure. Grey lines are WLC curves. **(C)** Refolding frequency, quantified as the fraction of relax-wait-stretch cycles showing core refolding, without (blue) and with (green) NAC (5 µM). N(cycles) are 193 (14 molecules) and 147 (17 molecules) without and with NAC. **** indicates significant difference (P<0.0001) from one-tailed two-proportion Z-test. **(D)** Unfolding forces of refolded structures without (blue) and with (green) NAC (5 µM), which show no significant difference (Mann-Whitney test), with close median values (23 pN without and 26 pN with 5 µM NAC). **(E)** Unfolding length of refolded core structures, without (blue) and with (green) NAC (5 µM), with mean values of 90.1 ± 5.8 nm (SD, N=98) and 91.1 ± 11.5 nm (SD, N=130), respectively. Histograms are normalized to the total number of states analyzed per condition.

To further understand NAĆs chaperoning mechanism, we analyzed the core states (**Figure 6B**, left panel) for their mechanical stability and unfolding step-size, as a proxy of their structural nature. No significant differences were observed between the average core unfolding forces in the absence and presence of NAC (**Figure 6D**), in line with MBP adopting the same or similar core states. Consistently, the measured length increase upon core unfolding peaked near the theoretically expected length of 96 nm for the core state, in both conditions (**Figure 6E**, methods). However, this core unfolding step-size distribution was narrower in the presence of NAC than in its absence, with the latter showing increased distribution in lower values down to about 70 nm. We quantified the relative abundance of these incomplete ‘small cores’ (between 70 and 90 nm) normalized by the total number of observed cores, and found that NAC caused a two-fold reduction (**Figure S11A**). Incomplete core structures are expected to be less stable mechanically, and we indeed found that their mean unfolding force was lower than for normal cores (**Figure S11B**). Hence, NAC not only promotes the formation of core states (**Figure 6C**), but those NAC-promoted core states are also more complete (**Figures 6E, S11A and S11B**).

The observed NAC chaperoning functions are notable. First, it is unclear for most chaperones if they promote folding directly, by lowering the folding energy barrier, or indirectely, by limiting aggregation interactions. As in this single-molecule experiment aggregation partners were absent, our data indicate that interactions with NAC are sufficient to directly promote folding transitions, as observed for the ATP-driven chaperone GroEL-ES^52^. Second, single-molecule and structural methods have shown that several chaperones including trigger factor, DnaK, and GroEL directly bind and stabilize partially folded intermediates, including the MBP core state^47–49^. Our data indicate that NAC promotes folding without stabilizing intermediate tertiary structures, given that NAC did not cause increased unfolding forces (**Figures 6D and S11B**). This NAC mechanism thus differs from that of other co-translationally acting chaperones, yet is consistent with early NAC binding to primary or secondary structure motifs as discussed above (**Figures 3I and 3J**), as these interactions do not increase tertiary structure stability. Thus, can directly promote native folding transitions without relying on tertiary structure stabilization.

## Discussion

NAC has an established role as molecular interaction hub that coordinates the access of protein biogenesis factors to nascent chains at the ribosomal tunnel exit^2–4,11–15,36^, but the nature and specificity of its interaction with NCs and its full impact on protein biogenesis remains unexplored. In this study, we fill this gap by profiling NAĆs interactions with ribosomes and nascent chains at the proteome level *in vivo*, defining its substrate pool and the nascent chain features recognized. Using single-molecule and cryo-EM approaches *in vitro*, we provide direct evidence of NAC’s dynamic interaction with emerging NCs, demonstrate its activity as chaperone and explore its working principles. These findings provide a conceptual framework for co-translational protein folding in eukaryotes, revealing a key role for NAC as a general chaperone for early folding steps at the tunnel exit.

Our SeRP data show that NAC is constitutively associated with ribosomes translating cytonuclear and mitochondrial proteins, with its β-barrel domain forming additional transient interactions with NCs during translation. Approximately 70 % of nascent cytonuclear and mitochondrial proteins experience at least one NAC binding period (**Table S1**). The binding periods correlate with the emergence of folded domains, suggesting a role for NAC as a chaperone. Our collective *in vivo* profiling, smFRET and cryo-EM experiments indicate that the NAC β-barrel domain dynamically engages NCs emerging at the tunnel exit, especially hydrophobic regions with α-helical propensity and amphipathic character, that are destined to form compact folded domains once all required residues become available (**Figure 3**). Through these local interactions, NAC favors early compaction of NCs (**Figure 5D**), which can promote folding. This model is supported by our optical tweezer experiments (**Figure 6C**), which show that NAC is capable of promoting native folding in polypeptide chains without stabilization of partial folding states. NAC also effectively suppress the presence of incompletely folded structures. This chaperone mechanism shows differences to known mechanisms such as shielding unfolded conformers as a ‘holdase’ (e.g. small HSPs, Hsp70), stabilizing folding intermediates (TF, Hsp70, Hsp90, GroEL-ES), and confining the folding of unfolded polypeptides (GroEL-ES)^47–49,53,54^.

We propose that several NAC features contribute to its chaperone activity. First, with an average binding period of 26 codons/residues (**Figure S3A**), NAC interactions are shorter than full domain lengths, indicating a role in guiding the early formation of small folding units. Supporting this, our cryo-EM data suggest that the RPL4 NC adopts a compact state at the tunnel exit upon NAC engagement, constrained by the available space between NAC and the ribosome vestibule. Additionally, NAC’s β-barrel domain interactions are dynamically regulated by NC length and features, with durations ranging from 5 to 25 seconds, as revealed by smFRET experiments (**Figure 4G**). We speculate that these context-dependent interactions facilitate NC compaction as it progressively emerges from the tunnel, creating a favorable microenvironment for early structure formation, such as helix-turn-helix elements (**Figure 7A**). Notably, NAC’s β-barrel domain presents both a hydrophobic pocket and a lysine-rich cationic surface. Together with the anionic ribosomal surface, this may form a multivalent environment for NC folding at the ribosomal vestibule. NAC may thus influence NC folding dynamics in multiple ways, depending on NC features, ribosome interactions, and domain size.

**Figure 7:**
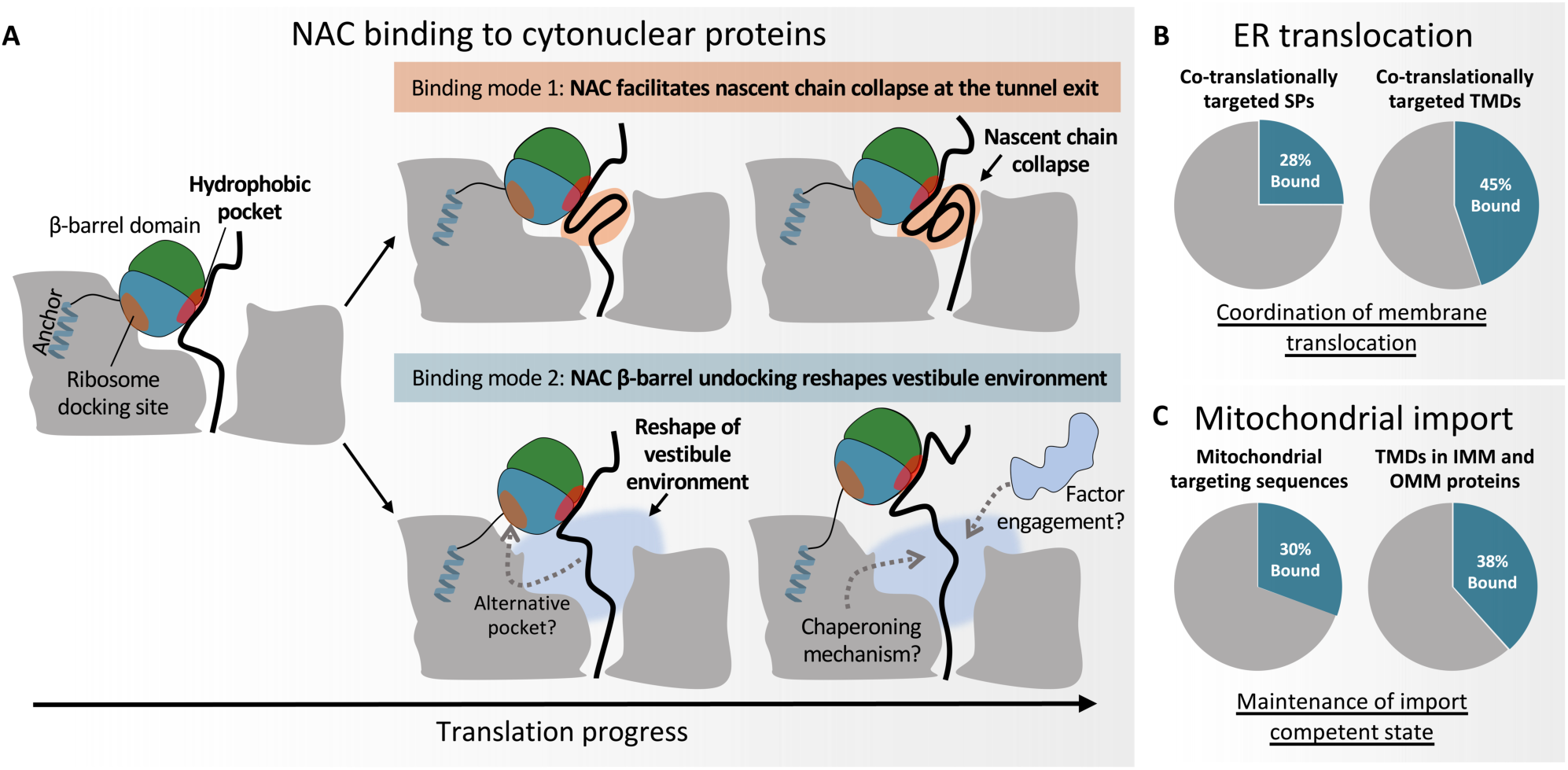
NAC roles in cytonuclear, mitochondrial and ER targeted proteins biogenesis. (**A**) Model of NAC chaperone action on cytonuclear proteins. NAC samples two distinct binding modes based on the position of the β-barrel domain. In binding mode 1, the NAC β-barrel stays ribosome-associated and facilitates NC collapse and early folding of the emerging polypeptide (orange). In binding mode 2, the NAC β-barrel dissociates from the ribosome upon binding the NC, thereby reshaping the vestibule environment (light blue), which could have diverse functional implications including increased accessibility to other factors. (**B**) NAC participates in ER membrane translocation by recognizing emerging SPs and, preferentially, TMDs. (**C**) NAC binding to certain MTS and TMDs in mitochondrial IMMs and OMMs, suggesting a general role in maintaining a translocation competent state of mitochondrial precursors, by either direct action on NC or coordination of downstream factors.

The dynamic changes in NAC interactions with the ribosome and the NC, as found for RPL4, could potentially be relevant to reshape the vestibule environment and allow NC handoff to other biogenesis factors (**Figure 7A**). This scenario may be functionally connected to NAC’s interplay with SRP; NAC regulates SRP through weakened binding of the β-barrel domain to the tunnel exit when signal peptides emerge, allowing SRP access to the NCs of ER-destined proteins^3^. Moreover, ribosome undocking of the β-barrel domain exposes a second, previously described pocket for SP binding in NAC^3^ (**Figure 7A**). Thus, by modulating its position and exposing an additional interaction site, NAC may act as a molecular switch that fine-tunes co-translational decisions, ensuring proper coordination between early folding events, membrane targeting and potentially other co-translational processes. Further studies will be needed to elucidate the broader functional implications of these dynamic interactions in shaping the protein biogenesis network.

Our proteome-wide profiling data identified the cellular nascent chain substrates of NAC. They include many ER-targeted proteins, with binding patterns that agree with the reported mechanism of NAC action in SRP-mediated co-translational ER targeting^3^ (**Figure 7B**). Additionally, we found NAC participates in mitochondrial membrane protein biogenesis by interacting with emerging TMDs of outer and inner membrane proteins (**Figure 7C**), which utilize distinct import pathways, presumably for transient shielding of these hydrophobic segments from aggregation. NAC also binds some MTS, the functional significance of which remains unclear. Certain MTS were recently found to co-translationally recruit Hsc70 and Hsp90 co-chaperones to maintain import competence^55^. NAC binding to specific MTS may be associated with MTS-mediated recruitment of such cytosolic factors that sustain import competence. Finally, NAC also associates with emerging domains of mitochondrial precursor proteins, which may enable domain-associated chaperoning activity for mitochondrial proteins, similar to its role in cytonuclear proteins (**Figure 3A and 3B**). Consequently, the mitochondrial import defects observed upon NAC depletion^16,38–40,42,56^ may arise from a failed chaperoning of soluble domains or a global disruption in polytopic membrane biogenesis, rather than a direct effect on specific import pathways. While we cannot rule out a role of NAC in mitochondrial targeting through direct binding or handover to other factors, our data suggest a scenario in which NAC influences mitochondrial import independently of the substratés translocation machinery.

In conclusion, NAC acts as a general coordinator of ribosome and NC accessibility while serving as a broad co-translational chaperone. These findings provide new insights into NAĆs multifaceted role in protein biogenesis.

## Acknowledgements

B.B., S.J.T. and N.B. acknowledge a research grant of the European Union (ERC - SyG - 101072047 - CoTransComplex). Views and opinions expressed are however those of the authors only and do not necessarily reflect those of the European Union or the European Research Council. Neither the European Union nor the granting authority can be held responsible for them. B.B. and G.K. are supported by the Deutsche Forschungsgemeinschaft (DFG, German Research Foundation; project number KR3593/4-1), S.S. and R.J.G. by grant R35 GM136321 from the National Institute of Health and grant 2219287 from the National Science Foundation, NB by grant (SNSF grant 310030_212308) from the Swiss National Science Foundation, J.S. by a longterm EMBO postdoctoral fellowship and M.P. by a Boehringer Ingelheim Fonds PhD fellowship. We thank the Klaus Tschira Foundation for their generous support in the acquisition of the sequencing device, and members of the Mayer, Ban, and Bukau labs for insightful discussions. We also thank B. Echeverria Perez for preparing the NAC construct used for single molecule experiments.

## Authors contributions

Conceptualization: J.S., M.G., R.J.G., M.I., M.P., N.B, S.T, S.S., G.K. and B.B. Methodology: J.S., M.G., R.J.G., M.I., M.P., I.E.K., N.B, S.T, S.S., G.K. and B.B. Investigation: J.S., M.G., R.J.G., M.I., M.P., I.E.K., P.D., D.C. and J.J. Software: J.S., M.G. and F.T. Formal analysis, Data curation, and Visualization: J.S., M.G., R.J.G., M.I., M.P., I.E.K., N.B, S.T, S.S., G.K. and B.B. Writing – original draft: J.S., G.K. and B.B. Writing – review & editing: all authors. Supervision and Funding acquisition: N.B, S.T, S.S., G.K. and B.B.

## Supplemental information

Document S1. Figures S1–S13, Tables S1 and S5

Table S2. Excel file containing additional data too large to fit in a PDF. High-confidence co-translationally ER-targeted substrates.

Table S3. Excel file containing additional data too large to fit in a PDF. Binding of NAC to mitochondrial MTS-containing proteins

Table S4. Excel file containing additional data too large to fit in a PDF.Binding of NAC to mitochondrial TMD-containing proteins.

## Materials and methods

### Cell culture

HEK293-T cells (Homo sapiens embryonal kidney, DSMZ Cat# ACC 635) were cultured in a humidified 37°C incubator with 5% CO_2_ using high glucose DMEM media containing GlutaMAX™ and pyruvate (Gibco) supplemented with 10% heat-inactivated FCS (Gibco), 100 U/mL penicillin and 100 μg/mL streptomycin (Gibco). Cells were seeded on 15 cm dishes 18–24 hours prior to harvest to ensure an 80–90% confluency was reached at the time of collection.

### Selective ribosome profiling

#### Purification of NAC-bound RNCs

NAC-bound RNCs were purified from a cell lysate or from sucrose cushion purified ribosomes after *in vivo* crosslinking to capture NC-independent (SeRP_Ribo_) or NC-dependent (SeRP_NC_) interactions, respectively.

For the SeRP_Ribo_ assay, HEK293-T cells were harvested from 2-3 plates by detachment with ice-cold 1x PBS supplemented with 100 µg/mL cycloheximide and 12 mM MgCl_2_, then pelleted by centrifugation. Cells were lysed in lysis buffer (20 mM Tris, pH 8.0, 140 mM KCl, 1 mM PMSF, 12 mM MgCl_2_, 0.5% NP-40, 100 μg/mL cycloheximide, 25 U/mL DNase1 (Roche), 1x protease inhibitor cocktail (complete EDTA free, Roche)). This experiment was performed with a lysis buffer containing either 70 mM (rep1) or 140 mM KCl (rep2) to account for potential salt sensitivity of NAC association. Both datasets were highly correlated, showing a lack of salt sensitivity within this concentration range (**Figure S12A**, R2 = 0.979). The lysate was cleared by centrifuging for 5 min at 20,000 *g* and 4°C, followed by five times trituration through a 23-G needle. The cleared lysate was digested with 550 U/mg RNA of RNase I (Ambion) at 4°C for 40 minutes after which the digest was stopped by adding Superase·In (Invitrogen). One part of the digestion reaction (following RNase I addition) was immediately subjected to immunoprecipitation (IP) by mixing with 70 μL of protein A Dynabeads (ThermoFisher), pre-coated with 10 μg of a homemade NACβ antibody, and incubated at 4°C for 40 minutes under rotation. NACβ was chosen as target for immunoprecipitation (IP) because this subunit contains the ribosomal anchor domain. The antigen-bound dynabeads were washed four times with 280 μL cold washing buffer (20 mM Tris, pH 8.0, 140 mM NaCl, 1 mM PMSF, 0.5% NP-40, 100 μg/mL cycloheximide, 12 mM MgCl_2_) and shock-frozen in liquid nitrogen. For isolation of TT samples, the remaining of the digestion reaction was pelleted by sucrose cushion centrifugation (25% (w/v) sucrose, 20 mM Tris, pH 8.0, 140 mM KCl, 12 mM MgCl_2_, 1 mM PMSF, 100 μg/mL cycloheximide) at 75,000 rpm for 90 min at 4°C (S120-AT2 rotor, Sorvall Discovery M120 SE Ultracentrifuge), and resuspended in lysis buffer.

For the SeRP_NC_ experiment, HEK293-T cells grown in 10-13 plates were harvested and subjected to *in vivo* crosslinking with 1 mM DSP (Thermo Fisher) prior to cell lysis as described in reference^57^. Samples were lysed and digested as previously described for the SeRP_Ribo_ and subjected to sucrose cushion purification at the same conditions as used for the SeRP_Ribo_ TT isolation. A fraction of the resuspended ribosomal pellets was set aside as the TT fraction, while the remaining sample was subjected to IP using either a homemade NACα or NACβ antibody. IP samples were mixed with 200 μL protein A dynabeads pre-coated with 18 µg of NACα-antibody (homemade) or 18 µg of NACβ-antibody (homemade), respectively, and incubated overnight at 4°C under rotation. The antigen-bound dynabeads were washed three times with 800 μL cold high-salt washing buffer (20 mM Tris, pH 8.0, 650 mM NaCl, 1 mM PMSF, 0.1% NP-40, 100 μg/mL cycloheximide, 12 mM MgCl_2_) and shock frozen in liquid nitrogen.

#### SolubleRP

Cells were harvested as previously as described above (SeRP_Ribo_ assay) from five 15 cm dishes per replicate. Cells pellets were resuspended in 1 mL ice-cold SolubleRP-lysis buffer (50 mM HEPES pH 7.0, 10 mM MgCl_2_, 140 mM KCl, 0.1 mg/mL cycloheximide, 0.02 U/μl DNaseI (Roche), 2 µg/mL E64, 5 µg/mL Leupeptin, 8 µg/mL Pepstatin and 40 µg/mL Bestatin). Resuspended cells were frozen in liquid nitrogen as droplets and stored at -80°C until lysis by mixer milling (2 min, 30 Hz, Retsch) under cryogenic conditions. Samples were thawed for 2 min at a 30°C water bath and RNA digestion was carried out using 650 U RNase I (Ambion) per mg of RNA for 30 min at 4°C. The lysate was then split into two halves to generate the soluble translatome and the TT. For the soluble translatome sample, the lysate was centrifuged twice at 16,000 g for 15 min, both times collecting the supernatant. For the total translatome sample, NP-40 was added to the lysate to a final concentration of 0.1%, incubated for 5 min on ice, and then centrifuged at 500 g for 5 min at 4°C. The supernatant from this sample, as well as the soluble lysate were subjected to sucrose cushion centrifugation and pellet fractions were resuspended in SolubleRP-lysis buffer.

Onset of membrane association was calculated by fitting ribosome depletion from the soluble fraction to a sigmoidal profile, using a sigmoidal fitting strategy as previously described^8^. High confidence co-translationally translocated substrates were annotated according to the following criteria: (i) An RPKM value of at least 10 in the TT, (ii) A depletion in the ST compared to the TT (ST/TT < 1), (iii) Soluble ribosome depletion follows a sigmoidal shape and the onset of assembly can be determined. To ensure accuracy, we performed manual curation to correct cases where the sigmoidal fitting introduced errors in the assignment.

#### Ribosome profiling library preparation and sequencing

Ribosome profiling libraries of SolubleRP and SeRP_NC_ samples were generated using classical ribosome profiling protocols as described previously^58^, in combination with custom rRNA depletion at the level of cDNA^8^. Ribosome profiling libraries of SeRP_Ribo_ were prepared using the NEXTflex Small RNA-Seq Kit v3 (PerkinElmer) following the manufacturer’s instructions. All libraries were sequenced on a NextSeq550 (Illumina) according to the manufacturer’s protocol and raw sequencing data was processed as described previously^8^, but keeping multimapped reads using star option ‘--outFilterMultimapNmax 20’.

Two independent replicates were prepared for each condition. ST replicates of each experiment were highly correlated (**Figure S12A**). The profiles obtained in the SeRP_NC_ experiments were virtually identical, independently of whether the IP was targeting NACα or NACβ (**Figure S12B**). We therefore selected the NACβ for further analysis, since this NAC subunit contains the ribosome anchor.

### Bioinformatic analysis

#### Single gene and metagene enrichment profiles

Single gene profiles show the position-wise 95% binomial confidence interval (CI) -according to Agresti and Coull-of the ST/TT enrichment corrected for library size. Annotated ribosome-exposed protein domains are shown by applying a 30-residue tunnel correction to account for their emergence from the exit tunnel. For metagene analysis, position-wise enrichments of ST over TT were calculated using a sliding window of 12 codons. Expression levels were used to normalize the contribution of each gene by dividing the read density at every position by the average read density of the gene in the TT. Genes with a coverage lower than 0.1 reads/codon were excluded. Metagene profiles were plotted as solid lines with a 95% confidence interval as previously described (https://github.com/ilia kats/RiboSeqTools and reference^8^). For both single and metagene plots, the two replicates of each experiment were merged into one dataset, increasing the read density and coverage. ST/TT enrichments were calculated as the ratio between the RPM of a given gene in the ST and TT.

#### Identification of NAC binding periods

To identify NAC binding periods, a 30-codon smoothing was applied to the ST and TT raw data to reduce experimental noise. Only proteins with a RPKM value higher than 10 in the TT were included in the analysis. Then, the 95% CI -according to Agresti and Coull-of the ST/TT enrichment was calculated for each individual replicate and used to define NAC-bound positions using a simple heuristic based on a threshold enrichment value of 1.5. Each position in a gene was assigned a value as follows:

1: High-confidence binding, if the lower CI was above 1.5.

0.2: Low-confidence binding, if the CI overlapped the 1.5 threshold.

0: Not bound, if the upper CI bound was below the threshold.

Positions with an average value across the two replicates higher that 0.6 were considered binding periods. Binding periods shorter than 10 consecutive codons were excluded. Proteins with at least one NAC binding period in the SeRP_NC_ were considered positive NAC substrates.

#### Annotation of the human proteome

Human protein domains were extracted from Manriquez-Sandoval et al.^59^, and analyzed based on the ECOD annotation of protein topologies and architectures. The DSSP annotation of the human proteome was performed using all human Alphafold derived .cif files (UP000005640_9606_HUMAN_v4) considering regions with a pLDDT value > 70. TMDs, SP and mitochondrial targeting sequences (MTS) were retrieved from UniprotKB^60^. TMDs associated with co-translational targeting were defined as the TMD emerging from the exit tunnel closer to the onset of membrane association. Mitochondrial protein location (matrix, IMP, OMP or intermembrane space) was extracted from MitoCarta3.0^61^.

Annotations of human protein subcellular localizations were extracted from the UniprotKB, Gene Ontology, and LOCATE^60,62,63^. If no information was present for the human protein, the localization of mouse and rat orthologues were instead retrieved. To identify cytonuclear proteins, the database was screened for the following terms: ‘cytosol’, ‘nucleoplasm’, ‘nucleus’, ‘cytoplasm’, ‘nucleoli’, ‘nucleolus’, ‘perinuclear region of cytoplasm’. Proteins were annotated as cytonuclear if they contained at least one of these terms and no TMD was annotated in UniprotKB. To avoid multiple annotations for the same protein, annotations were hierarchically ranked as following: (i) Annotations from different organisms were ranked human > mouse > rat; (ii) Annotations from different databases within the same organism were ranked UniprotKB > Gene Ontology > LOCATE.

#### Analysis of sequence features

Residue hydrophobicity was calculated using Kyte-Doolittle hydrophobicity scale^64^. α-helical hydrophobic moment (μH) of protein sequences was used as a measure of amphiphilicity of α-helices and was calculated as described by Eisenberg and co-workers^65^ using a sliding window of 11 residues. α-helical and β-sheet content was calculated based on the previously mentioned DSSP annotation derived from Alphafold derived structures of human proteins^66,67^. Protein disorder was calculated with IUPRED 2.0 using default settings^68^. For the metagene analysis of protein features aligned to the onset of NAC binding periods position-wise mean confidence intervals were calculated for each feature. Background profiles were generated by randomly sampling 5 positions in each of the analyzed genes.

#### Gene ontology and frequency enrichments

Gene ontology terms were analyzed with the Functional Annotation Tool of DAVID v6.8 (Database for Annotation, Visualization and Integrated Discovery)^69^. The analysis was performed with default settings, and custom datasets were uploaded as background sets tailored to the tested enrichment. GO terms with an adjusted p-value < 0.05 were considered significantly enriched. Simple frequency enrichments were calculated for domain topology and architecture by calculating fold enrichment, and statistical significance was tested using Fisher’s exact test.

#### Position-specific enrichment of DSSP secondary structure and amino-acid identity

DSSP secondary structure assignment and amino acid identity of the ribosome-proximal 60 residues at the onset of NAC binding periods were extracted and compared with a random background generated by randomly sampling 5 positions in each of the analyzed genes. Enrichment analysis were performed using Two Sample Logo webpage^70^ using default settings.

### Protein Preparation for smTIRF microscopy

#### NAC Purification

The human NAC heterodimer, comprising N-terminally 6×His tagged NACα and NACβ with the S57C mutation for labeling, was expressed using a pET28b plasmid in *E. coli* BL21 (DE3) cells, as described^35^. Cells were grown at 37°C to OD_600_ 0.6, at which the temperature was reduced to 18°C, and overexpression was induced with 1 mM IPTG for 16 hours.

Harvested cells were lysed by sonication in NAC Lysis Buffer (50 mM KHEPES, pH 7.5; 1 M NaCl; 6 mM β-mercaptoethanol; 30 mM imidazole; 10% glycerol) supplemented with 1×ProBlock Gold Protease Inhibitor Cocktail (GoldBio) and 1 mM AEBSF. The lysate was clarified by centrifugation at 18,000 rpm for 1 hour in a JA-20 rotor (Beckman Coulter). The clarified lysate was incubated with Ni Sepharose High-Performance resin (Cytiva) at a ratio of 1 ml resin per liter of culture for 1 hour with gentle rotation at 4°C. The resin was subsequently washed in batches with 50 column volumes (CVs) of NAC Lysis Buffer. Bound proteins were eluted twice with 5 CVs of NAC Elution Buffer (50 mM HEPES-KOH, pH 7.5; 100 mM NaCl; 500 mM imidazole; 1 mM TCEP; 10% glycerol). Eluted protein was incubated with PreScission Protease overnight at 4 °C with gentle rotation. Protein was further purified on a Mono Q 10/100 GL anion-exchange column (Cytiva) in NAC Buffer (50 mM HEPES-KOH, pH 7.5; 1 mM TCEP; 10% glycerol) using a linear gradient of 100 mM – 500 mM NaCl over 10 column volumes. Peak fractions were pooled and concentrated using a 10 kDa molecular weight cutoff (MWCO) centrifugal concentrator (Amicon). The purified protein was aliquoted, flash-frozen in liquid nitrogen, and stored at −80 °C. NAC concentration was determined using an extinction coefficient of ε_280_ = 2,980 M⁻¹ cm⁻¹.

#### NAC labeling

NACα/NACβ(S57C) was labeled with maleimide-Cy3B (Cytiva). The protein was exchanged into labeling buffer (50 mM KHEPES, pH 7.5; 100 mM NaCl; 1 mM TCEP; 20% glycerol) and incubated with a 5-fold molar excess of dye for 2 hours at room temperature. Excess free dye was removed using a G-25 Sephadex size exclusion column (GE Healthcare). Fractions containing labeled NAC were identified via SDS-PAGE, pooled, and concentrated to a final concentration of 150 µM.

### RNC_hRPL4_ Purification

#### PCR and in vitro transcription

DNA fragments encoding, from 5’ to 3’, the T7 promoter, IRES, 3X-FLAG, SUMO, and the nascent chain at defined lengths were generated by PCR. All amplicons contained an additional CAC at the 3’ end to generate a C-terminal Val that stabilizes the stalled ribosome-nascent chain complex. Amber codons were inserted in the RPL4 coding sequence to allow for non-natural amino acid incorporation and dye labeling. PCR products were purified using the QIAquick PCR Purification Kit (Qiagen, cat no. 28104) and used for *in vitro* transcription using home-made T7 polymerase and the MegaScript protocol. 5 mM 5’-biotin-GMP (TriLink) was included during transcription to incorporate a biotin at the 5’end of the mRNA. mRNA was partially purified by precipitation in 3M LiCl at –20 °C for an hour, followed by wash with 200 uL of ice-cold 70% ethanol. RNA pellet was resuspended in Ultrapure d.d. water.

#### RNC labeling and purification

RNCs were generated by *in vitro* translation in rabbit reticulocyte lysate (RRL, Green Hectares) supplemented with 1 µM *Methanosarcina mazei* pyrrolysine synthase (*Mm*PylRS), 10 mg/L *M. mazei* amber suppressor tRNA (*Mm*PyltRNA), and 100 µM axial-trans-cyclooct-2-en-L-lysine (TCOK, SiChem), for 30 minutes at 32 °C, as described^35^. Protocols for *Mm*PylRS and *Mm*PyltRNA preparation were described previously^35^.

Translation reactions were layered onto a high-salt sucrose cushion (50 mM KHEPES, pH 7.5, 1 M KOAc, 15 mM Mg(OAc)₂, 0.5 M sucrose, 0.1% Triton X-100, 2 mM DTT) at a 2:3 volumetric ratio and ultracentrifuged at 100,000 rpm for 45 minutes at 4 °C (TLA100.3 rotor, Beckman Coulter). Ribosomal pellets were resuspended in RNC buffer (50 mM KHEPES, pH 7.5, 150 mM KOAc, 2 mM Mg(OAc)₂) to a ribosome concentration of 1 µM, and were incubated with 1 µM tetrazine-conjugated Atto647N dye (Jena Biosciences) for 30 minutes at 25 °C to enable Diels-Alder cycloaddition of the fluorescent dye to TCOK.

Labeled RNCs were incubated with anti-DYKDDDK magnetic agarose beads (Pierce) pre-equilibrated in RNC Buffer for one hour at 4 °C with constant rotation. Beads were washed sequentially with 10 CVs of RNC buffer containing 300 mM KOAc, 10 CVs of RNC buffer with 0.1% Triton X-100, and 10 CVs of RNC buffer alone. RNCs were eluted using 1.0 mg/mL 3×FLAG peptide (DYKDDDK) for 30 minutes at 4 °C with constant rotation. The eluate was treated with 1 µM 3C protease (GoldBio) at 25 °C for at least one hour. Purified RNCs were sedimented through a high-salt sucrose cushion at a 2:3 volumetric ratio by ultracentrifugation at 100,000 rpm for 30 minutes at 4 °C (TLA120.2 rotor, Beckman Coulter). The pellet was resuspended in Assay Buffer (50 mM KHEPES, pH 7.5, 150 mM KOAc, 5 mM Mg(OAc)₂, 0.04% Nikkol, 2 mM DTT) to a final ribosome concentration of ∼100 nM. Fluorescently labeled nascent chains were validated by SDS-PAGE and fluorescence imaging (Typhoon biomolecular imager, Cytiva). Labeled RNCs were snap-frozen in liquid nitrogen and stored at −80 °C.

### Single-molecule TIRF microscopy

Quartz imaging slides and coverslips were aminosilanized using Vectabond (Vector Laboratories) and coated with biotin-PEG (Laysan Bio). The slides were passivated in Passivation Buffer (1× Tween-20, 1 mg/mL BSA) for at least 1 hour at 25 °C, washed with 50 mM KHEPES, pH 7.5, and incubated with 0.5 mg/mL NeutrAvidin (ThermoFisher) for 10 minutes. Excess NeutrAvidin was removed by washing with Imaging Buffer, comprising the Assay Buffer supplemented with 1 mg/mL BSA and the oxygen scavenging system^71^ (4 mM Trolox, 2.5 mM protocatechuic acid (PCA), and 50 nM protocatechuate-3,4-dioxygenase (PCD, Sigma)).

Biotinylated RNCs were diluted to 1.5 nM in Imaging Buffer and immobilized on NeutrAvidin-functionalized slides for 10 minutes at 25 °C. Following that, the slides were flushed with 2 nM Cy3B-labeled NAC in Imaging Buffer. Images were captured using MicroManager on a custom-built (TIRF) microscopy setup^72^. The presence of Atto647N-labeled RNCs was verified by excitation at 635 nm. Single-molecule fluorescence measurements were conducted with donor excitation at 532 nm (Cy3B) in single-excitation mode, with simultaneous detection of donor and acceptor channels at a temporal resolution of 50 ms.

### Single-molecule FRET data analysis

Donor and acceptor channels were aligned and analyzed using iSMS software^73^. Raw donor and acceptor fluorescence intensities were corrected for background noise, the γ factor that accounts for differences in dye quantum yield, and donor leakage into the acceptor channel. Apparent FRET efficiencies (*E_app_*) were calculated using Equation 1,

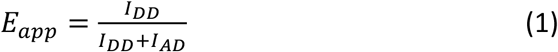

where *I*_*DD*_ and *I*_*AD*_ are the corrected fluorescence intensities of the donor and acceptor dyes, respectively, upon excitation of the donor.

The FRET traces from all movies for the same experimental group were pooled and analyzed by Hidden Markov Modeling (HMM) using the hmmlearn algorithm available in Python. The number of FRET states was determined using the Bayesian Information Criterion (BIC), which identified three distinct FRET populations. The mean FRET efficiencies for the individual states, along with its standard deviation, were used to fit smFRET histograms to a Gaussian mixture model using Equation 2,

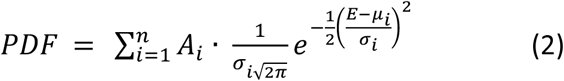

where PDF is the probability density function, *n* is the number of FRET states detected and was set to 3 (determined by HMM), *A*_*i*_ is the weight of the i^th^ Gaussian, and σ_*i*_ and μ_*i*_ are the standard deviation and center (determined by HMM) of the i^th^ Gaussian.

To evaluate the residence time of NAC on RNCs, the cumulative fluorescence intensity (*I*_cumulative_) for the dye pairs in experiment 2 was calculated for each trace using Equation 3:

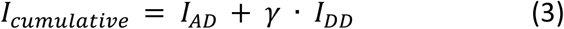

Colocalization of labeled NAC and RNCs was identified when *I*_cumulative_ exceeded background noise. Two-state HMM was used to distinguish bound and dissociated states of NAC, allowing the determination of dwell times (*t*) for individual colocalization events. The cumulative probability distribution of *t* was fit to the two-exponential decay function in Equation 4,

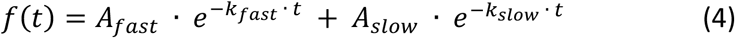

in which *A_fast_* and *A_slow_* are the amplitudes of the fast- and slow-dissociating NAC populations, and *k_fast_* and *k_slow_* are their respective dissociation rate constants.

### Cryo-EM sample preparation

#### Design of RPL4(1-158)-Xbp1u plasmid

The plasmid encoded a T7 promoter, EMCV internal ribosome entry site, EGFP, SENP-EuB protease cleavage site, the first 158 amino acids of human RPL4, Xbp1u translational arrest peptide and a 30-nt polyA tail, in the pT7CFE1-Chis vector (Thermo Fischer Scientific).

Due to partial folding of the 26 amino acid-long Xbp1u peptide in the polypeptide exit tunnel^45^, its length is approximately equivalent to that of a 22 amino acid unfolded polypeptide chain. Therefore, the nascent chain (after GFP cleavage) mimics 180 amino acid-long RPL4, assuming that the C-terminal 30 amino acid stretch of RPL4 in the polypeptide exit tunnel is unfolded.

#### *In vitro* transcription of mRNA [RPL4(1-158)-Xbp1u]

Linearized DNA template for *in vitro* transcription, which encompassed the T7 promoter at the 5’ end and the polyA tail at 3’ end, was generated by PCR using the Q5 DNA polymerase (NEB). After treating the PCR reaction with DpnI (NEB), the template was purified using the QIAquick PCR purification kit (Qiagen). The *in vitro* transcription reaction consisted of 1.15 mg/ml homemade T7 polymerase, 12 nM DNA template, 5 mM NTPs, 30 mM MgCl_2_, 40 mM Tris-HCl pH 7.6, 1 mM DTT, 2 mM spermidine, and 0.1 U/µL RNaseOUT^TM^ Recombinant Ribonuclease Inhibitor (Thermo Fischer Scientific). The reaction was carried out at 37 °C for 2 h, followed by digestion of the DNA template by 0.04 U/µL of RNase-free DNase I (Qiagen). mRNA was precipitated by ½ reaction volume of 7M LiCl at -20 °C for 1 h and pelleted by centrifugation for 15 min at 13,000 rpm and 4 °C. The pellet was washed with 70% ethanol and centrifuged again, then air-dried. The mRNA was resuspended in pure water and stored at -20 °C.

#### Protein purification

The human NAC heterodimer, containing NACα with an N-terminal 6x His tag and TEV protease cleavage site, was expressed in *E. coli* BL21 (DE3). Transformed bacteria were grown in 2YT containing kanamycin (50 µg/mL) at 37 °C to an optical density (OD_600_) of 0.7, followed by induction with 1 mM isopropyl β-D-1-thiogalactopyranoside (IPTG) and incubation overnight at 18 °C. Bacteria were lysed by sonication in buffer A (50 mM Hepes pH 7.6, 800 mM KCl, 40 mM imidazole, 10% glycerol), supplemented with 100 μM PMSF, 2 μM E−64, 10 μM Leupeptin, 10 μM Bestatin, 1 μM Pepstatin A, 10 μg/mL Aprotinin, and a pinch of DNase I. The lysate was centrifuged in an SS-34 rotor (Sorvall) for 45 min at 20,000 rpm and 4 °C. The clarified supernatant was loaded on a HisTrap HP column (Cytiva), washed with buffer A, and eluted directly on a HiTrap Q HP column (Cytiva) using a linear gradient from buffer A to 100% buffer B (50 mM Hepes pH 7.6, 100 mM KCl, 500 mM imidazole, 10% glycerol) over 10 column volumes. The protein was then washed with buffer C (50 mM Hepes pH 7.6, 100 mM KCl, 10% glycerol), and eluted with a linear gradient from buffer C to 100% buffer D (50 mM Hepes pH 7.6, 800 mM KCl, 10% glycerol) over 10 column volumes. Peak fractions were pooled and incubated with homemade TEV protease overnight at 4°C. The protein was loaded on a HisTrap HP column and washed with buffer A, followed by a linear gradient from buffer A to 100% buffer B over 10 column volumes. NAC was collected in the flowthrough, concentrated and buffer exchanged to 50 mM Hepes pH 7.6, 150 mM KOAc, 10% glycerol, 1 mM TCEP, using a 30 K MWCO centrifugal filter (Amicon). Aliquots were flash-frozen in liquid nitrogen and stored at -80°C. GADD34 was purified as described in reference^74^.

#### Generation and purification of RNC (RPL4(1-158)-Xbp1u)

Rpl4 RNC were generated using the human *in vitro* translation system as described previously^74^. The *in vitro* translation reaction consisted of 75% (v/v) human cell extract, 0.01 mg/ml GADD34, 100 nM mRNA, 15 mM Hepes pH 7.0, 120 mM KCl, 0.9 mM MgCl_2_ and 20 mM creatine phosphatase. The reaction was incubated at 33 °C for 20 min.

RPL4 RNC were purified using GFP nanobody-conjugated magnetic beads and elution by GFP cleavage (tag off) as described previously^75^ with the following modifications: The *in vitro* translation reaction was supplemented with 0.1% Triton X-100 and diluted to 1.2 mL with buffer 1T (50 mM Hepes pH 7.4, 100 mM KCl, 5 mM MgCl_2_, 0.1% Triton X-100), followed by incubation with the GFP nanobody-conjugated magnetic beads for 1 h at 4 °C with rotation. Beads were washed twice with buffer 1T, twice with buffer 2T (50 mM Hepes pH 7.4, 500 mM KCl, 5 mM MgCl_2_, 0.1% Triton X-100), and twice with buffer 1T. RNCs were eluted with 250 nM SENP-EuB SUMO protease (homemade) in buffer 1T by incubating for 20 min on ice; the elution was performed twice. The pooled elutions were clarified at 21,300 *g* for 1 min, followed by centrifugation in the TLA-55 rotor (Beckman Coulter) at 55,000 rpm for 2 h. The RNC pellet was resuspended in RNC buffer (50 mM Hepes pH 7.4, 100 mM KOAc, 5 mM MgOAc_2_). Aliquots were flash-frozen in liquid nitrogen and stored at -80 °C.

Samples were taken throughout RNC generation and purification for analysis on 4-12% SDS PAGE gels (Millipore), followed by transfer using the eBlot^TM^ L1 system (Genscript) and detection using a polyclonal GFP primary antibody (Invitrogen, A6455) and HRP-conjugated mouse anti-rabbit secondary antibody (Santa Cruz Biotechnology, sc-2357).

#### Reconstitution of RNC/NAC complex for cryo-EM

Prior to grid preparation, 100 nM RPL4 RNC was mixed with 1 µM NAC heterodimer and 0.02% octaethylene glycol monododecyl ether in RNC buffer. The mixture was incubated at 30 °C for 10 min, then placed on ice.

### Cryo-EM methods

#### Grid preparation

Quantifoil R2/2 Cu 300 grids, coated with a homemade 1 nm thick layer of amorphous carbon, were glow-discharged for 15 s at 15 mA using the PELCO easiGlow Discharge cleaning system (Ted Pella). Grids were mounted into a Vitrobot MK IV (Thermo Fischer Scientific), preset to 4 °C and 100% humidity, and 5 μl of the RNC/NAC sample was applied on the grid. After 30 s, the grids were blotted for 4 s with a blot force of 15, then immediately plunge-frozen in a 1:2 mixture of ethane and propane. Grids were imaged using a Titan Krios G3i transmission electron microscope at 300 kV, equipped with a BioQuantum imaging filter with the energy filter slit set to 20 e^-^V, and a K3 direct electron detector (Gatan). Movies were collected in super-resolution counting mode, then binned to the physical pixel size during data collection. The nominal magnification was 81000x, corresponding to a physical pixel size of 1.06 Å/pixel. During collection, exposure was set to reach a total electron dose of 50 e^-^/Å ^2^ and the defocus values ranged from -0.7 µm to -2.2 µm in steps of 0.3 µm. EPU 3.6.0.3689 (Thermo Fischer Scientific) was used for automated data collection. 9,751 and 10,444 movies were collected in two independent sessions from two different RNC and cryo-EM grid preparations.

#### Cryo-EM data processing

CryoSPARC Live was used to motion-correct, dose-weigh and sum movie frames into micrographs, followed by estimation of CTF parameters (**Figure S13**). Some micrographs were excluded based on the maximum resolution of the CTF fit, the ice thickness and total motion. Particles were picked either with an 80S template (low-pass filtered to 40 Å) in CryoSPARC or using a blob picker (circular, 250 - 350 Å) in CryoSPARC Live. Particles were extracted using a box size of 560 pixels and subsequently Fourier cropped to a box size of 186 pixels. All subsequent processing was performed in CryoSPARC^76^ for both datasets. After 2D classification, ribosome-like classes were selected for further processing (371,689 particles in dataset 1; 503,918 particles in dataset 2). An *ab initio* reconstructed 80S ribosome was used as a template for heterogeneous refinement, which separated a class of leading ribosomes with a P-site tRNA and no downstream ribosome (207,513 particles in dataset 1; 233,286 particles in dataset 2). As these represent RNCs that are translationally arrested at the programmed site and bear the designed nascent chain, these particles were selected for further processing and homogenously refined. The particles were re-extracted with a box size of 560 pixels and homogenously refined, with optimization of per-particle defocus and per-group CTF parameters. Due to the high similarity of cryo-EM maps from the two datasets, particles were merged at this point and homogenously refined. 3D classification with a mask around NAC was used to select particles with a well-defined NAC globular domain (104,588 particles; 24% of leading ribosomes). These were homogenously refined with per-particle defocus optimization, and subsequently non-uniformly refined to an overall resolution of 2.8 Å. The map was further locally refined with a mask covering the 40S for the purposes of model building only.

#### Model building

For molecular interpretation of the 2.8 Å EM map calculated for the NACα/β-bound 80S ribosome stalled on the RPL4-XBP1u mRNA construct, available structures for the human small ribosomal subunit (8PPL), the large subunit (9GMO), the tRNA(Met)-bound Xbp1u nascent chain (from GR7Q) and a model of human NACα/β (from 7QWR) were docked as rigid bodies using CHIMERAX^77^. The composite model was the manually readjusted in COOT^78^ using available high-resolution structures of human (8QOI) and rabbit (7O7Y) 80S ribosomes as a guide.

The yeast tRNA(Met) sequence was altered to the human (available from the GtRNAdb on https://gtrnadb.ucsc.edu/)^79^. The C-terminal portion of the nascent chain could be CA-traced from residues 151–184, with unambiguous assignment of protein side chains from residues 156–171 (**Figure 5C**) and from Ala179 to the tRNA-attached Met184:

MACARPLISVYSEKGESSGKNVTLPAVFKAPIRPDIVNFVHTNLRKNNRQPYAVSELAGHQTSAESWGTGRAVARIPRVRGGGTHRSGQGAFGNMCRGGRMFAPTKTWRRWHRRVNTTQKRYAICSALAASALPALVMSKGHRIEEV PELPLVVEDKVQKDPVPYQPPFLCQWGRHQCAWKPL**M**N

Protein sequence of the nascent chain construct, with the N-terminal portion corresponding to RPL4 residues 1–158 and the C-terminally fused XBP1 stalling sequence underlined. The traced portion (residues 151–184) is marked in grey, and the tRNA(Met)-attached methionine in bold.

This, together with careful inspection of the codon-anticodon base pairing and the nucleotide pattern of the tRNA(Met) anticodon stem-loop in a map focused on the small subunit, confirmed that the translation had been stalled by the Xbp1 sequence at the expected position.

#### Real space refinement

The completed structure was refined for several cycles into the 2.8 Å EM map using real space refinement in PHENIX version 1.21.2-5419^80^. Custom restraints for modified rRNA and protein residues were generated using phenix.eLBOW^80^, PRODRG^81^ or exported from COOT^78^. To stabilize good model geometry also in less well-ordered areas at the periphery of the 80S map, the model was initially refined with added hydrogens and using protein secondary structure, rotamer and Ramachandran as well as RNA base pair and stacking restraints. Real space difference maps were used to find and correct remaining discrepancies between the model and the EM map. During final cycles of refinement, hydrogens were removed from the model, and the secondary structure restraints were released except for the Ramachandran restraints for a better model-to-map fit. The structure was validated using MOLPROBITY^82^, revealing that the model displays excellent geometry and map correlation (**Table S5**). The resolution of the refined model vs. map FSC at a value of 0.5 (3.0 Å, masked) coincides well with the resolution determined between the map half-sets at the FSC=0.143 criterion (2.8 Å).

### Single-molecule optical tweezers

Bacterial MBP with introduced cysteine residues was expressed in BL21 and affinity-purified using Amylose resin (New England Biolabs)^46^. Human NAC heterodimer, comprising NACα and hist-tagged NACβ, was co-expressed in BL21 and purified by Ni-NTA and MonoQ chromatography^3^. MBP was coupled to maleimide-modified 20-nt DNA oligos (biomers.net) via thiol-maleimide reaction, and hybridized to 1.3-kbp double-stranded functionalized DNA handles via an overhang approach^46^. This yielded a DNA-MBP-DNA construct, functionalized with biotin group at one end and digoxigenin at the opposite end. To create the first bead-construct connection, the construct was incubated with anti-digoxigenin antibody-coated polystyrene beads (diameter 2.1 μm, Roche) for 30 mins with rotary shaking at 4 °C in assay buffer (50 mM HEPES, 150 mM potassium acetate, 1 mM TCEP, pH 7.4). The second bead-construct connection was created by using neutravidin-coated polystyrene beads (diameter 2.1 μm, Spherotech). Single-molecule measurements were performed in the assay buffer with addition of an oxygen scavenging system (3 units mL^-1^ pyranose oxidase, 90 units mL^-1^ catalase, and 50 mM glucose, all from Merck).

All single-molecule optical trapping measurements were performed using a C-Trap setup (Lumicks)^46,49^, with a constant trapping laser intensity. In these measurements, the samples were introduced into separate channels of a pressure-controlled microfluidic flow cell. Following optical trapping of a bead pair, which contained an anti-digoxigenin-bead coupled to the MBP construct and a neutravidin-bead, the two beads were moved close to enable tether formation, verified by increase in force upon retraction. Single tethers were confirmed by the typical shape of the MBP unfolding profile with a large increase in extension due to unfolding of the MBP core and by the presence of the DNA overstretching plateau at 65 pN. The tethers were repeatedly stretched and relaxed, with the pulling and relaxation rates of the steerable trap of 0.1 µms^-1^ and 5 s waiting time at 0 pN, until tether breakage. For each molecule, the optical traps were calibrated by fitting a Lorenzian function to the power spectrum of the trapped bead pair.

Data were collected at 50 Hz and decimated to 500 Hz for the analysis. Segments of the force-extension curves were fit with two worm-like chain (WLC) models in series: an extensible WLC for the DNA contribution, and the Odijk approximation for an inextensible WLC for the protein component^46,49^, with rulers at 0 nm (fully folded MBP), 33 nm (folded core) and 129 nm (fully unfolded protein). First pulling curves were excluded from the analysis, as we aimed to study refolding from the unfolded state. Any tethers that did not show full unfolding in the first pulling cycle were excluded, as well as any molecules that did not display any core unfolding transitions.

## References

1. Cassaignau, A.M.E., Cabrita, L.D., and Christodoulou, J. (2020). How Does the Ribosome Fold the Proteome? Annu Rev Biochem 89, 389–415. 10.1146/annurev-biochem-062917-012226.

2. Gamerdinger, M., Jia, M., Schloemer, R., Rabl, L., Jaskolowski, M., Khakzar, K.M., Ulusoy, Z., Wallisch, A., Jomaa, A., Hunaeus, G., et al. (2023). NAC controls cotranslational N-terminal methionine excision in eukaryotes. Science 380, 1238–1243. 10.1126/science.adg3297.

3. Jomaa, A., Gamerdinger, M., Hsieh, H.-H., Wallisch, A., Chandrasekaran, V., Ulusoy, Z., Scaiola, A., Hegde, R.S., Shan, S.-O., Ban, N., et al. (2022). Mechanism of signal sequence handover from NAC to SRP on ribosomes during ER-protein targeting. Science 375, 839–844. 10.1126/science.abl6459.

4. Lentzsch, A.M., Yudin, D., Gamerdinger, M., Chandrasekar, S., Rabl, L., Scaiola, A., Deuerling, E., Ban, N., and Shan, S.-O. (2024). NAC guides a ribosomal multienzyme complex for nascent protein processing. Nature 633, 718–724. 10.1038/s41586-024-07846-7.

5. Gilmore, R., Blobel, G., and Walter, P. (1982). Protein translocation across the endoplasmic reticulum. I. Detection in the microsomal membrane of a receptor for the signal recognition particle. J Cell Biol 95, 463–469. 10.1083/jcb.95.2.463.

6. Halic, M., Blau, M., Becker, T., Mielke, T., Pool, M.R., Wild, K., Sinning, I., and Beckmann, R. (2006). Following the signal sequence from ribosomal tunnel exit to signal recognition particle. Nature 444, 507–511. 10.1038/nature05326.

7. Streit, J.O., Bukvin, I.V., Chan, S.H.S., Bashir, S., Woodburn, L.F., Włodarski, T., Figueiredo, A.M., Jurkeviciute, G., Sidhu, H.K., Hornby, C.R., et al. (2024). The ribosome lowers the entropic penalty of protein folding. Nature 633, 232–239. 10.1038/s41586-024-07784-4.

8. Bertolini, M., Fenzl, K., Kats, I., Wruck, F., Tippmann, F., Schmitt, J., Auburger, J.J., Tans, S., Bukau, B., and Kramer, G. (2021). Interactions between nascent proteins translated by adjacent ribosomes drive homomer assembly. Science 371, 57–64. 10.1126/science.abc7151.

9. Shiber, A., Döring, K., Friedrich, U., Klann, K., Merker, D., Zedan, M., Tippmann, F., Kramer, G., and Bukau, B. (2018). Cotranslational assembly of protein complexes in eukaryotes revealed by ribosome profiling. Nature 561, 268–272. 10.1038/s41586-018-0462-y.

10. Shieh, Y.-W., Minguez, P., Bork, P., Auburger, J.J., Guilbride, D.L., Kramer, G., and Bukau, B. (2015). Operon structure and cotranslational subunit association direct protein assembly in bacteria. Science 350, 678–680. 10.1126/science.aac8171.

11. Gamerdinger, M., Kobayashi, K., Wallisch, A., Kreft, S.G., Sailer, C., Schlömer, R., Sachs, N., Jomaa, A., Stengel, F., Ban, N., et al. (2019). Early Scanning of Nascent Polypeptides inside the Ribosomal Tunnel by NAC. Mol Cell 75, 996–1006.e8. 10.1016/j.molcel.2019.06.030.

12. Wiedmann, B., Sakai, H., Davis, T.A., and Wiedmann, M. (1994). A protein complex required for signal-sequence-specific sorting and translocation. Nature 370, 434–440. 10.1038/370434a0.

13. Klein, M., Wild, K., and Sinning, I. (2024). Multi-protein assemblies orchestrate co-translational enzymatic processing on the human ribosome. Nat Commun 15, 7681. 10.1038/s41467-024-51964-9.

14. Minoia, M., Quintana-Cordero, J., Jetzinger, K., Kotan, I.E., Turnbull, K.J., Ciccarelli, M., Masser, A.E., Liebers, D., Gouarin, E., Czech, M., et al. (2024). Chp1 is a dedicated chaperone at the ribosome that safeguards eEF1A biogenesis. Nat Commun 15, 1382. 10.1038/s41467-024-45645-w.

15. Gamerdinger, M., Hanebuth, M.A., Frickey, T., and Deuerling, E. (2015). The principle of antagonism ensures protein targeting specificity at the endoplasmic reticulum. Science 348, 201–207. 10.1126/science.aaa5335.

16. Fünfschilling, U., and Rospert, S. (1999). Nascent polypeptide-associated complex stimulates protein import into yeast mitochondria. Mol Biol Cell 10, 3289–3299. 10.1091/mbc.10.10.3289.

17. Raue, U., Oellerer, S., and Rospert, S. (2007). Association of protein biogenesis factors at the yeast ribosomal tunnel exit is affected by the translational status and nascent polypeptide sequence. J Biol Chem 282, 7809–7816. 10.1074/jbc.M611436200.

18. Wegrzyn, R.D., Hofmann, D., Merz, F., Nikolay, R., Rauch, T., Graf, C., and Deuerling, E. (2006). A conserved motif is prerequisite for the interaction of NAC with ribosomal protein L23 and nascent chains. J Biol Chem 281, 2847–2857. 10.1074/jbc.M511420200.

19. Deuerling, E., Gamerdinger, M., and Kreft, S.G. (2019). Chaperone Interactions at the Ribosome. Cold Spring Harb Perspect Biol 11, a033977. 10.1101/cshperspect.a033977.

20. Wang, F., Durfee, L.A., and Huibregtse, J.M. (2013). A cotranslational ubiquitination pathway for quality control of misfolded proteins. Mol Cell 50, 368–378. 10.1016/j.molcel.2013.03.009.

21. Duttler, S., Pechmann, S., and Frydman, J. (2013). Principles of cotranslational ubiquitination and quality control at the ribosome. Mol Cell 50, 379–393. 10.1016/j.molcel.2013.03.010.

22. Deuerling, E., and Bukau, B. (2004). Chaperone-assisted folding of newly synthesized proteins in the cytosol. Crit Rev Biochem Mol Biol 39, 261–277. 10.1080/10409230490892496.

23. Preissler, S., and Deuerling, E. (2012). Ribosome-associated chaperones as key players in proteostasis. Trends Biochem Sci 37, 274–283. 10.1016/j.tibs.2012.03.002.

24. Kirstein-Miles, J., Scior, A., Deuerling, E., and Morimoto, R.I. (2013). The nascent polypeptide-associated complex is a key regulator of proteostasis. EMBO J 32, 1451–1468. 10.1038/emboj.2013.87.

25. Shen, K., Gamerdinger, M., Chan, R., Gense, K., Martin, E.M., Sachs, N., Knight, P.D., Schlömer, R., Calabrese, A.N., Stewart, K.L., et al. (2019). Dual Role of Ribosome-Binding Domain of NAC as a Potent Suppressor of Protein Aggregation and Aging-Related Proteinopathies. Mol Cell 74, 729–741.e7. 10.1016/j.molcel.2019.03.012.

26. Martin, E.M., Jackson, M.P., Gamerdinger, M., Gense, K., Karamonos, T.K., Humes, J.R., Deuerling, E., Ashcroft, A.E., and Radford, S.E. (2018). Conformational flexibility within the nascent polypeptide-associated complex enables its interactions with structurally diverse client proteins. J Biol Chem 293, 8554–8568. 10.1074/jbc.RA117.001568.

27. Koplin, A., Preissler, S., Ilina, Y., Koch, M., Scior, A., Erhardt, M., and Deuerling, E. (2010). A dual function for chaperones SSB-RAC and the NAC nascent polypeptide-associated complex on ribosomes. J Cell Biol 189, 57–68. 10.1083/jcb.200910074.

28. Zhu, Z., Wang, S., and Shan, S.-O. (2022). Ribosome profiling reveals multiple roles of SecA in cotranslational protein export. Nat Commun 13, 3393. 10.1038/s41467-022-31061-5.

29. Stein, K.C., Kriel, A., and Frydman, J. (2019). Nascent Polypeptide Domain Topology and Elongation Rate Direct the Cotranslational Hierarchy of Hsp70 and TRiC/CCT. Mol Cell 75, 1117–1130.e5. 10.1016/j.molcel.2019.06.036.

30. Galmozzi, C.V., Merker, D., Friedrich, U.A., Döring, K., and Kramer, G. (2019). Selective ribosome profiling to study interactions of translating ribosomes in yeast. Nat Protoc 14, 2279– 2317. 10.1038/s41596-019-0185-z.

31. Döring, K., Ahmed, N., Riemer, T., Suresh, H.G., Vainshtein, Y., Habich, M., Riemer, J., Mayer, M.P., O’Brien, E.P., Kramer, G., et al. (2017). Profiling Ssb-Nascent Chain Interactions Reveals Principles of Hsp70-Assisted Folding. Cell 170, 298–311.e20. 10.1016/j.cell.2017.06.038.

32. Chartron, J.W., Hunt, K.C.L., and Frydman, J. (2016). Cotranslational signal-independent SRP preloading during membrane targeting. Nature 536, 224–228. 10.1038/nature19309.

33. Schibich, D., Gloge, F., Pöhner, I., Björkholm, P., Wade, R.C., von Heijne, G., Bukau, B., and Kramer, G. (2016). Global profiling of SRP interaction with nascent polypeptides. Nature 536, 219–223. 10.1038/nature19070.

34. Oh, E., Becker, A.H., Sandikci, A., Huber, D., Chaba, R., Gloge, F., Nichols, R.J., Typas, A., Gross, C.A., Kramer, G., et al. (2011). Selective ribosome profiling reveals the cotranslational chaperone action of trigger factor in vivo. Cell 147, 1295–1308. 10.1016/j.cell.2011.10.044.

35. Hsieh, H.-H., Lee, J.H., Chandrasekar, S., and Shan, S.-O. (2020). A ribosome-associated chaperone enables substrate triage in a cotranslational protein targeting complex. Nat Commun 11, 5840. 10.1038/s41467-020-19548-5.

36. del Alamo, M., Hogan, D.J., Pechmann, S., Albanese, V., Brown, P.O., and Frydman, J. (2011). Defining the specificity of cotranslationally acting chaperones by systematic analysis of mRNAs associated with ribosome-nascent chain complexes. PLoS Biol 9, e1001100. 10.1371/journal.pbio.1001100.

37. Becker, A.H., Oh, E., Weissman, J.S., Kramer, G., and Bukau, B. (2013). Selective ribosome profiling as a tool for studying the interaction of chaperones and targeting factors with nascent polypeptide chains and ribosomes. Nat Protoc 8, 2212–2239. 10.1038/nprot.2013.133.

38. Ponce-Rojas, J.C., Avendaño-Monsalve, M.C., Yañez-Falcón, A.R., Jaimes-Miranda, F., Garay, E., Torres-Quiroz, F., DeLuna, A., and Funes, S. (2017). αβ’-NAC cooperates with Sam37 to mediate early stages of mitochondrial protein import. FEBS J 284, 814–830. 10.1111/febs.14024.

39. Lesnik, C., Cohen, Y., Atir-Lande, A., Schuldiner, M., and Arava, Y. (2014). OM14 is a mitochondrial receptor for cytosolic ribosomes that supports co-translational import into mitochondria. Nat Commun 5, 5711. 10.1038/ncomms6711.

40. George, R., Beddoe, T., Landl, K., and Lithgow, T. (1998). The yeast nascent polypeptide-associated complex initiates protein targeting to mitochondria in vivo. Proc Natl Acad Sci U S A 95, 2296–2301. 10.1073/pnas.95.5.2296.

41. Wiedemann, N., and Pfanner, N. (2017). Mitochondrial Machineries for Protein Import and Assembly. Annu Rev Biochem 86, 685–714. 10.1146/annurev-biochem-060815-014352.

42. Muthukumar, G., Stevens, T.A., Inglis, A.J., Esantsi, T.K., Saunders, R.A., Schulte, F., Voorhees, R.M., Guna, A., and Weissman, J.S. (2024). Triaging of α-helical proteins to the mitochondrial outer membrane by distinct chaperone machinery based on substrate topology. Mol Cell 84, 1101–1119.e9. 10.1016/j.molcel.2024.01.028.

43. Pillet, B., García-Gómez, J.J., Pausch, P., Falquet, L., Bange, G., de la Cruz, J., and Kressler, D. (2015). The Dedicated Chaperone Acl4 Escorts Ribosomal Protein Rpl4 to Its Nuclear Pre-60S Assembly Site. PLoS Genet 11, e1005565. 10.1371/journal.pgen.1005565.

44. Lin, Z., Gasic, I., Chandrasekaran, V., Peters, N., Shao, S., Mitchison, T.J., and Hegde, R.S. (2020). TTC5 mediates autoregulation of tubulin via mRNA degradation. Science 367, 100–104. 10.1126/science.aaz4352.

45. Shanmuganathan, V., Schiller, N., Magoulopoulou, A., Cheng, J., Braunger, K., Cymer, F., Berninghausen, O., Beatrix, B., Kohno, K., von Heijne, G., et al. (2019). Structural and mutational analysis of the ribosome-arresting human XBP1u. Elife 8, e46267. 10.7554/eLife.46267.

46. Avellaneda, M.J., Koers, E.J., Minde, D.P., Sunderlikova, V., and Tans, S.J. (2020). Simultaneous sensing and imaging of individual biomolecular complexes enabled by modular DNA-protein coupling. Commun Chem 3, 20. 10.1038/s42004-020-0267-4.

47. Mashaghi, A., Kramer, G., Bechtluft, P., Zachmann-Brand, B., Driessen, A.J.M., Bukau, B., and Tans, S.J. (2013). Reshaping of the conformational search of a protein by the chaperone trigger factor. Nature 500, 98–101. 10.1038/nature12293.

48. Mashaghi, A., Bezrukavnikov, S., Minde, D.P., Wentink, A.S., Kityk, R., Zachmann-Brand, B., Mayer, M.P., Kramer, G., Bukau, B., and Tans, S.J. (2016). Alternative modes of client binding enable functional plasticity of Hsp70. Nature 539, 448–451. 10.1038/nature20137.

49. Naqvi, M.M., Avellaneda, M.J., Roth, A., Koers, E.J., Roland, A., Sunderlikova, V., Kramer, G., Rye, H.S., and Tans, S.J. (2022). Protein chain collapse modulation and folding stimulation by GroEL-ES. Sci Adv 8, eabl6293. 10.1126/sciadv.abl6293.

50. Bechtluft, P., van Leeuwen, R.G.H., Tyreman, M., Tomkiewicz, D., Nouwen, N., Tepper, H.L., Driessen, A.J.M., and Tans, S.J. (2007). Direct observation of chaperone-induced changes in a protein folding pathway. Science 318, 1458–1461. 10.1126/science.1144972.

51. Dobson, C.M. (2003). Protein folding and misfolding. Nature 426, 884–890. 10.1038/nature02261.

52. Sharma, S., Chakraborty, K., Müller, B.K., Astola, N., Tang, Y.-C., Lamb, D.C., Hayer-Hartl, M., and Hartl, F.U. (2008). Monitoring protein conformation along the pathway of chaperonin-assisted folding. Cell 133, 142–153. 10.1016/j.cell.2008.01.048.

53. Mogk, A., Ruger-Herreros, C., and Bukau, B. (2019). Cellular Functions and Mechanisms of Action of Small Heat Shock Proteins. Annu Rev Microbiol 73, 89–110. 10.1146/annurev-micro-020518-115515.

54. Karagöz, G.E., and Rüdiger, S.G.D. (2015). Hsp90 interaction with clients. Trends Biochem Sci 40, 117–125. 10.1016/j.tibs.2014.12.002.

55. Juszkiewicz, S., Peak-Chew, S.-Y., and Hegde, R.S. (2025). Mechanism of chaperone recruitment and retention on mitochondrial precursors. Mol Biol Cell, mbcE25010035. 10.1091/mbc.E25-01-0035.

56. Ramalho, S., Alkan, F., Prekovic, S., Jastrzebski, K., Barberà, E.P., Hoekman, L., Altelaar, M., de Heus, C., Liv, N., Rodríguez-Colman, M.J., et al. (2025). NAC regulates metabolism and cell fate in intestinal stem cells. Sci Adv 11, eadn9750. 10.1126/sciadv.adn9750.

57. Günnigmann, M., Koubek, J., Kramer, G., and Bukau, B. (2023). Selective ribosome profiling as a tool to study interactions of translating ribosomes in mammalian cells. Methods Enzymol 684, 1–38. 10.1016/bs.mie.2022.09.006.

58. McGlincy, N.J., and Ingolia, N.T. (2017). Transcriptome-wide measurement of translation by ribosome profiling. Methods 126, 112–129. 10.1016/j.ymeth.2017.05.028.

59. Manriquez-Sandoval, E., and Fried, S.D. (2022). DomainMapper: Accurate domain structure annotation including those with non-contiguous topologies. Protein Sci 31, e4465. 10.1002/pro.4465.

60. UniProt Consortium (2019). UniProt: a worldwide hub of protein knowledge. Nucleic Acids Res 47, D506–D515. 10.1093/nar/gky1049.

61. Rath, S., Sharma, R., Gupta, R., Ast, T., Chan, C., Durham, T.J., Goodman, R.P., Grabarek, Z., Haas, M.E., Hung, W.H.W., et al. (2021). MitoCarta3.0: an updated mitochondrial proteome now with sub-organelle localization and pathway annotations. Nucleic Acids Res 49, D1541–D1547. 10.1093/nar/gkaa1011.

62. Sprenger, J., Lynn Fink, J., Karunaratne, S., Hanson, K., Hamilton, N.A., and Teasdale, R.D. (2008). LOCATE: a mammalian protein subcellular localization database. Nucleic Acids Res 36, D230–233. 10.1093/nar/gkm950.

63. Ashburner, M., Ball, C.A., Blake, J.A., Botstein, D., Butler, H., Cherry, J.M., Davis, A.P., Dolinski, K., Dwight, S.S., Eppig, J.T., et al. (2000). Gene ontology: tool for the unification of biology. The Gene Ontology Consortium. Nat Genet 25, 25–29. 10.1038/75556.

64. Kyte, J., and Doolittle, R.F. (1982). A simple method for displaying the hydropathic character of a protein. J Mol Biol 157, 105–132. 10.1016/0022-2836(82)90515-0.

65. Eisenberg, D., Weiss, R.M., and Terwilliger, T.C. (1982). The helical hydrophobic moment: a measure of the amphiphilicity of a helix. Nature 299, 371–374. 10.1038/299371a0.

66. Varadi, M., Bertoni, D., Magana, P., Paramval, U., Pidruchna, I., Radhakrishnan, M., Tsenkov, M., Nair, S., Mirdita, M., Yeo, J., et al. (2024). AlphaFold Protein Structure Database in 2024: providing structure coverage for over 214 million protein sequences. Nucleic Acids Res 52, D368–D375. 10.1093/nar/gkad1011.

67. Jumper, J., Evans, R., Pritzel, A., Green, T., Figurnov, M., Ronneberger, O., Tunyasuvunakool, K., Bates, R., Žídek, A., Potapenko, A., et al. (2021). Highly accurate protein structure prediction with AlphaFold. Nature 596, 583–589. 10.1038/s41586-021-03819-2.

68. Mészáros, B., Erdos, G., and Dosztányi, Z. (2018). IUPred2A: context-dependent prediction of protein disorder as a function of redox state and protein binding. Nucleic Acids Res 46, W329–W337. 10.1093/nar/gky384.

69. Huang, D.W., Sherman, B.T., and Lempicki, R.A. (2009). Systematic and integrative analysis of large gene lists using DAVID bioinformatics resources. Nat Protoc 4, 44–57. 10.1038/nprot.2008.211.

70. Vacic, V., Iakoucheva, L.M., and Radivojac, P. (2006). Two Sample Logo: a graphical representation of the differences between two sets of sequence alignments. Bioinformatics 22, 1536–1537. 10.1093/bioinformatics/btl151.

71. Aitken, C.E., Marshall, R.A., and Puglisi, J.D. (2008). An oxygen scavenging system for improvement of dye stability in single-molecule fluorescence experiments. Biophys J 94, 1826– 1835. 10.1529/biophysj.107.117689.

72. Shen, K., Arslan, S., Akopian, D., Ha, T., and Shan, S. (2012). Activated GTPase movement on an RNA scaffold drives co-translational protein targeting. Nature 492, 271–275. 10.1038/nature11726.

73. Preus, S., Noer, S.L., Hildebrandt, L.L., Gudnason, D., and Birkedal, V. (2015). iSMS: single-molecule FRET microscopy software. Nat Methods 12, 593–594. 10.1038/nmeth.3435.

74. Bothe, A., and Ban, N. (2024). A highly optimized human in vitro translation system. Cell Rep Methods 4, 100755. 10.1016/j.crmeth.2024.100755.

75. Stevens, T.A., Tomaleri, G.P., Hazu, M., Wei, S., Nguyen, V.N., DeKalb, C., Voorhees, R.M., and Pleiner, T. (2024). A nanobody-based strategy for rapid and scalable purification of human protein complexes. Nat Protoc 19, 127–158. 10.1038/s41596-023-00904-w.

76. Punjani, A., Rubinstein, J.L., Fleet, D.J., and Brubaker, M.A. (2017). cryoSPARC: algorithms for rapid unsupervised cryo-EM structure determination. Nat Methods 14, 290–296. 10.1038/nmeth.4169.

77. Pettersen, E.F., Goddard, T.D., Huang, C.C., Meng, E.C., Couch, G.S., Croll, T.I., Morris, J.H., and Ferrin, T.E. (2021). UCSF ChimeraX: Structure visualization for researchers, educators, and developers. Protein Sci 30, 70–82. 10.1002/pro.3943.

78. Emsley, P., Lohkamp, B., Scott, W.G., and Cowtan, K. (2010). Features and development of Coot. Acta Crystallogr D Biol Crystallogr 66, 486–501. 10.1107/S0907444910007493.

79. Chan, P.P., and Lowe, T.M. (2016). GtRNAdb 2.0: an expanded database of transfer RNA genes identified in complete and draft genomes. Nucleic Acids Res 44, D184–189. 10.1093/nar/gkv1309.

80. Liebschner, D., Afonine, P.V., Baker, M.L., Bunkóczi, G., Chen, V.B., Croll, T.I., Hintze, B., Hung, L.W., Jain, S., McCoy, A.J., et al. (2019). Macromolecular structure determination using X-rays, neutrons and electrons: recent developments in Phenix. Acta Crystallogr D Struct Biol 75, 861– 877. 10.1107/S2059798319011471.

81. Schüttelkopf, A.W., and van Aalten, D.M.F. (2004). PRODRG: a tool for high-throughput crystallography of protein-ligand complexes. Acta Crystallogr D Biol Crystallogr 60, 1355–1363. 10.1107/S0907444904011679.

82. Chen, V.B., Arendall, W.B., Headd, J.J., Keedy, D.A., Immormino, R.M., Kapral, G.J., Murray, L.W., Richardson, J.S., and Richardson, D.C. (2010). MolProbity: all-atom structure validation for macromolecular crystallography. Acta Crystallogr D Biol Crystallogr 66, 12–21. 10.1107/S0907444909042073.

