## Supplementary Materials for "NAC promotes co-translational protein folding at the ribosomal tunnel exit"

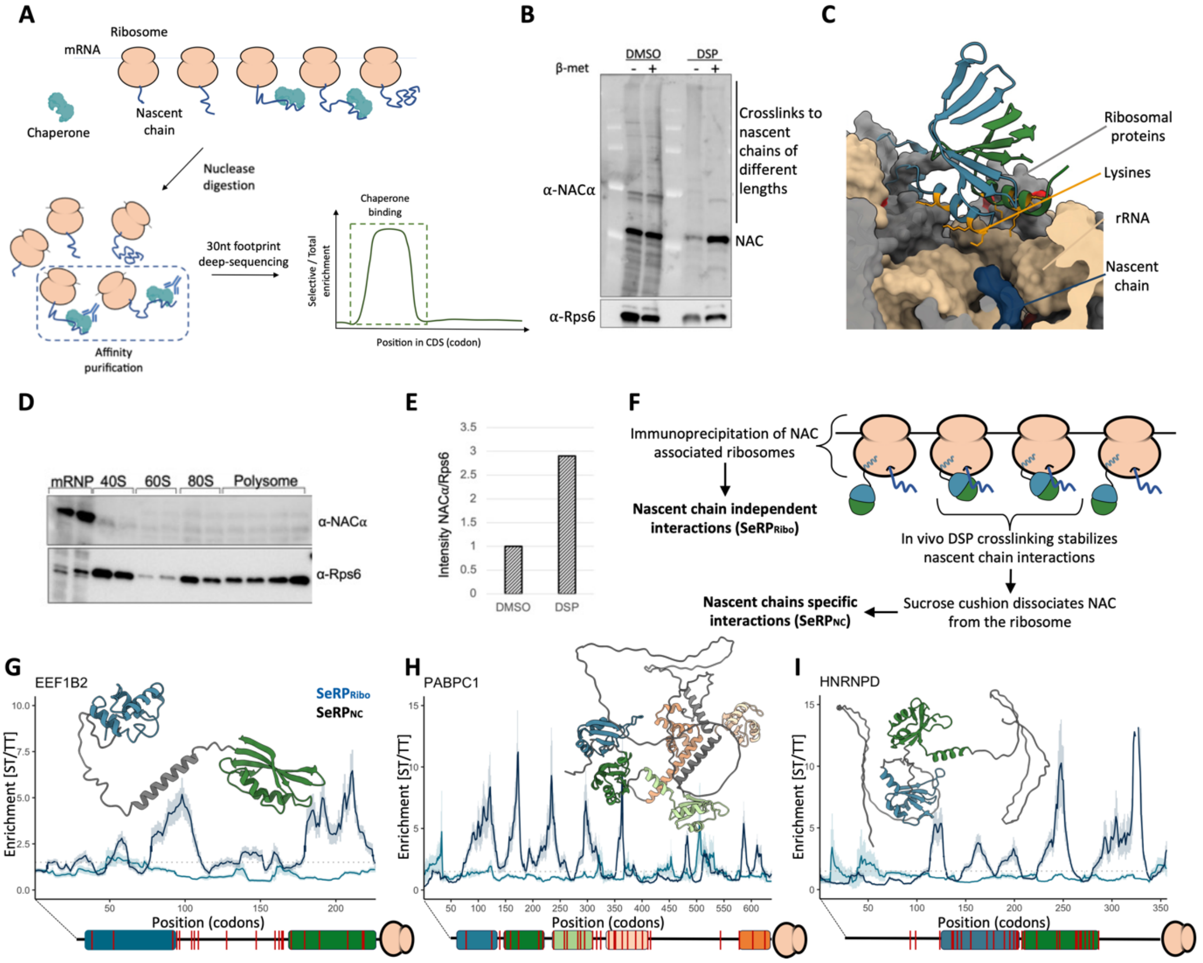


**Figure S1. Development of NAC-selective Ribosome Profiling (SeRP).** (**A**) Schematic representation of the SeRP pipeline. (**B**) Western blot using NAC- and ribosomal Rps6 (as control)-specific antibodies showing NAC association with ribosomes in untreated or *in vivo* DSP-crosslinked samples in absence and presence of β-mercaptoethanol. Ribosomes were purified by sucrose cushion centrifugation. (**C**) β-barrel domain of NAC harboring the ribosome binding site. rRNA and ribosomal proteins are colored in light brown and gray, respectively. Lysines of the ß-barrel domain contacting ribosomal RNA are colored in orange while lysines in ribosomal proteins are shown in red, being depleted near this binding site. (**D**) In the absence of crosslinker, NAC dissociates from ribosomes during sucrose gradient centrifugation. mRNP refers to the upper fractions of the gradient. (**E**) Relative quantification of NAC association with ribosomes from panel B. NAC-associated intensity was normalized using Rps6 as control (in presence of β-mercaptoethanol). (**F**) Schematic representation of the SeRP profiling strategies designed to assess the NAC co-translational interactome. (**G-I**) Nascent protein enrichment profiles of cytonuclear EEF1B2, PABPC1 and HNRNPD. NC-independent ($SeRP_{Ribo}$) and NC-dependent ($SeRP_{NC}$) enrichments are colored in blue and black, respectively. Domains (rectangles) and lysines (red bars) are annotated using a 30-residue tunnel correction to account for emergence from the exit tunnel, as depicted in panel A. Shown structures were extracted from Alphafold [1,2]. Dotted line indicates an enrichment of 1.5, used as a threshold to define binding periods. Shadowed areas indicate 95% confidence interval.

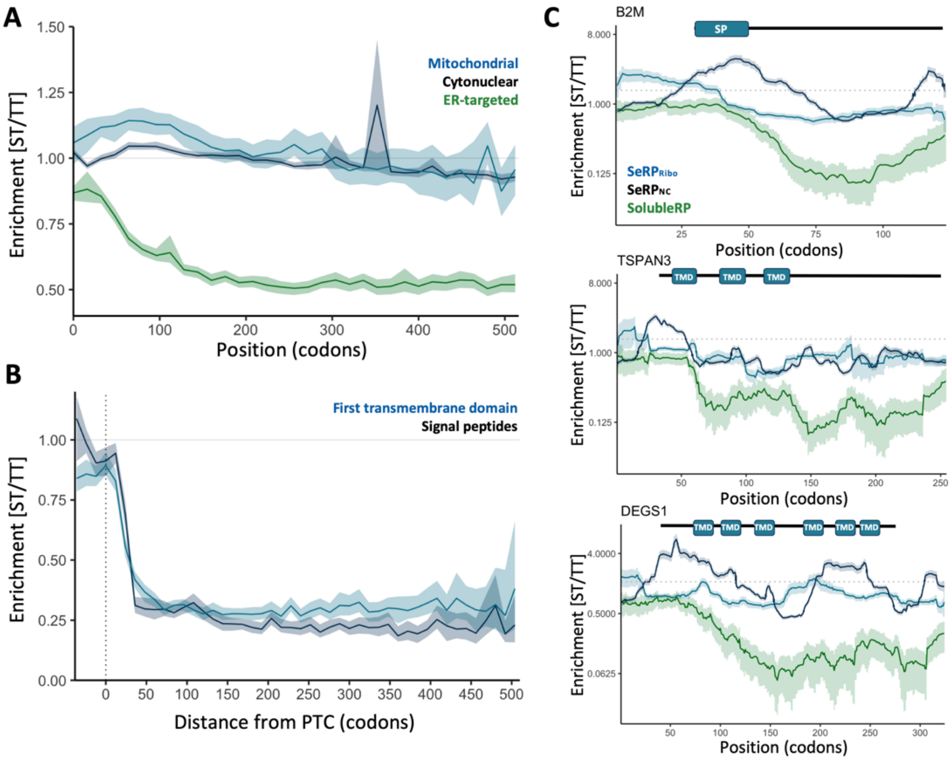


**Figure S2: Co-translational membrane association patterns in SolubleRP datasets.** (**A**) Metagene ST/TT enrichments of SolubleRP datasets based on cellular location. Shadowed areas indicate 95% confidence intervals. (**B**) SolubleRP ST/TT enrichment aligned to the synthesis of the last (C-terminal) residue in a signal peptide or the first (N-terminal) TMD in the protein sequence. At position 0, such feature is in the PTC. Shadowed areas indicate 95% confidence intervals. (**C**) Single nascent protein enrichment profiles of B2M, TSPAN3 and DEGS1. NC-dependent, NC-independent and SolubleRP enrichments are colored in blue, black and green, respectively. TMDs and SPs are annotated using a 30-residue tunnel correction to account for emergence from the tunnel. Shadowed areas indicate 95% confidence interval.

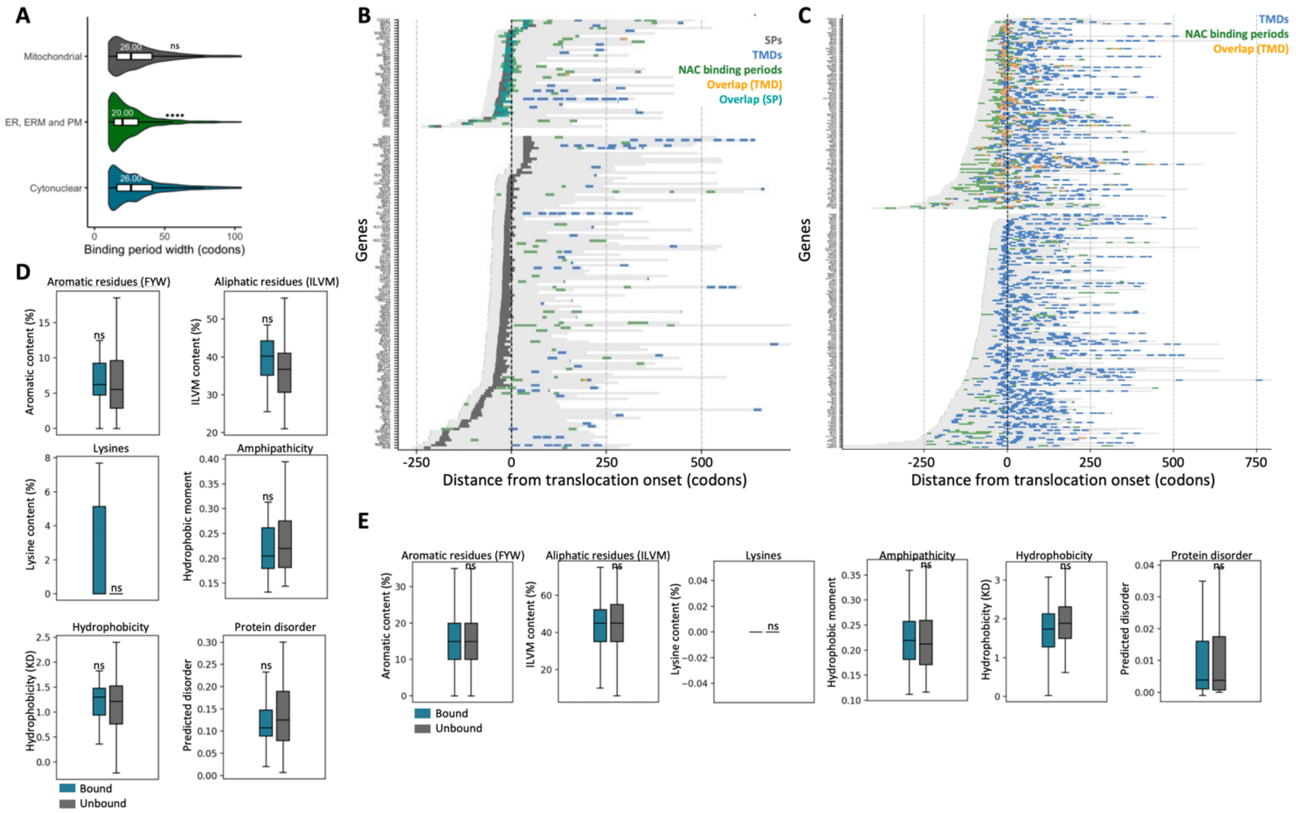


**Figure S3: Features of NAC binding to ER-targeted proteins.** (**A**) Violin plots showing the average length of NAC binding periods according to protein's cellular destiny. 12023, 1087 and 782 binding periods were analyzed for cytonuclear, ER-targeted and mitochondrial classes, respectively. Statistical significance was tested using a two-sided Mann-Whitney U test. ****: p-value < 0.0001; ns: not statistically significant. (**B**) Heat map of SP containing proteins in high confidence co-translationally translocated proteins aligned to the onset of membrane association. SPs, TMDs and NAC binding periods are indicated in gray, blue and green, respectively. Overlaps between NAC binding periods and SP or TMDs are colored in orange. SP and TMD positions have been corrected to account for their emergence from the exit tunnel (30 residue correction). Genes with overlap between NAC binding periods and SPs/TMDs are shown at the top. (**C**) Heat map of TMD containing proteins without SPs in high confidence co-translationally translocated proteins aligned to the onset of membrane association. TMDs, NAC binding periods and overlaps are indicated using the same color code than in panel D and the 30-residue tunnel correction. Genes with overlap between NAC binding periods and SPs/TMDs are shown at the top. (**D-E**) Comparison of features of SPs (D) and TMDs (E) bound and not bound by NAC. Only TMDs and SPs associated with co-translational translocation were included in the analysis. Statistical significance was tested using a two-sided Mann-Whitney U test. ns: not statistically significant

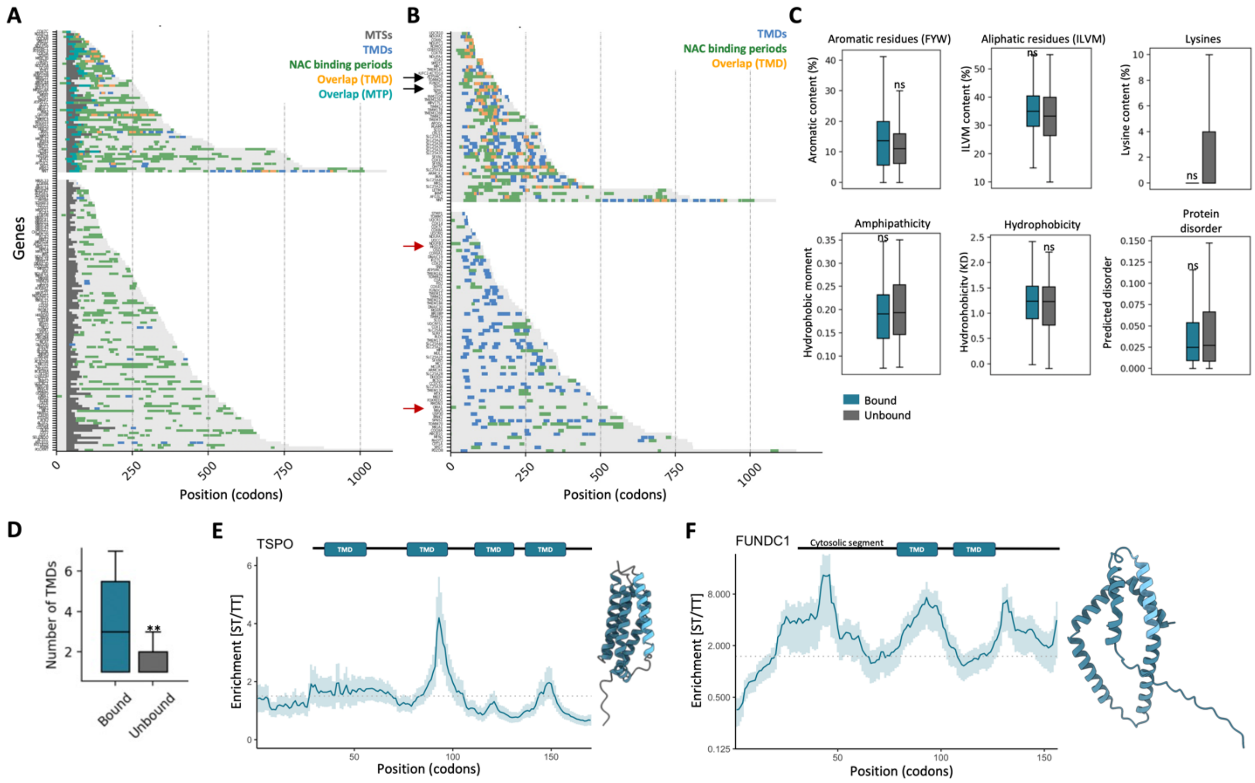


**Figure S4: NAC binding to MTSs and TMDs in mitochondrial proteins.** (**A**) Heat map of mitochondrial targeting sequences (MTSs) containing proteins. MTSs, transmembrane domains (TMDs) and NAC binding periods are indicated in gray, blue and green, respectively. Overlaps between NAC binding periods and SPs or TMDs are colored in bright blue and orange respectively. MTS and TMD positions have been corrected to account for their emergence from the exit tunnel (30-residue correction). Genes with overlap between NAC binding periods and MTS are shown at the top. (**B**) Heat map of TMD containing mitochondrial proteins without MTSs. TMDs, NAC binding periods and overlaps are indicated using the same color code as in panel A and the 30-residue tunnel correction. Genes with overlap between NAC binding periods and TMDs are shown at the top. (**C**) Features of TMDs bound and not bound by NAC based on the SeRP$_{NC}$ dataset. Statistical significance was tested using a two-sided Mann-Whitney U test. ns: not statistically significant. (**D**) Box-plot showing the number of TMDs in proteins with TMDs bound by NAC compared to those with unbound TMDs from the heat map in panel B. (**E-F**) Single nascent protein enrichment profiles of TSPO (E) and FUNDC1 (F) NC-dependent interactions. TMDs are annotated using a 30-residue tunnel correction to account for emergence from the tunnel. Alphafold predicted structures are shown, with the bound TMD in light blue. Shadowed areas indicate 95% Agresti-Coull confidence interval.

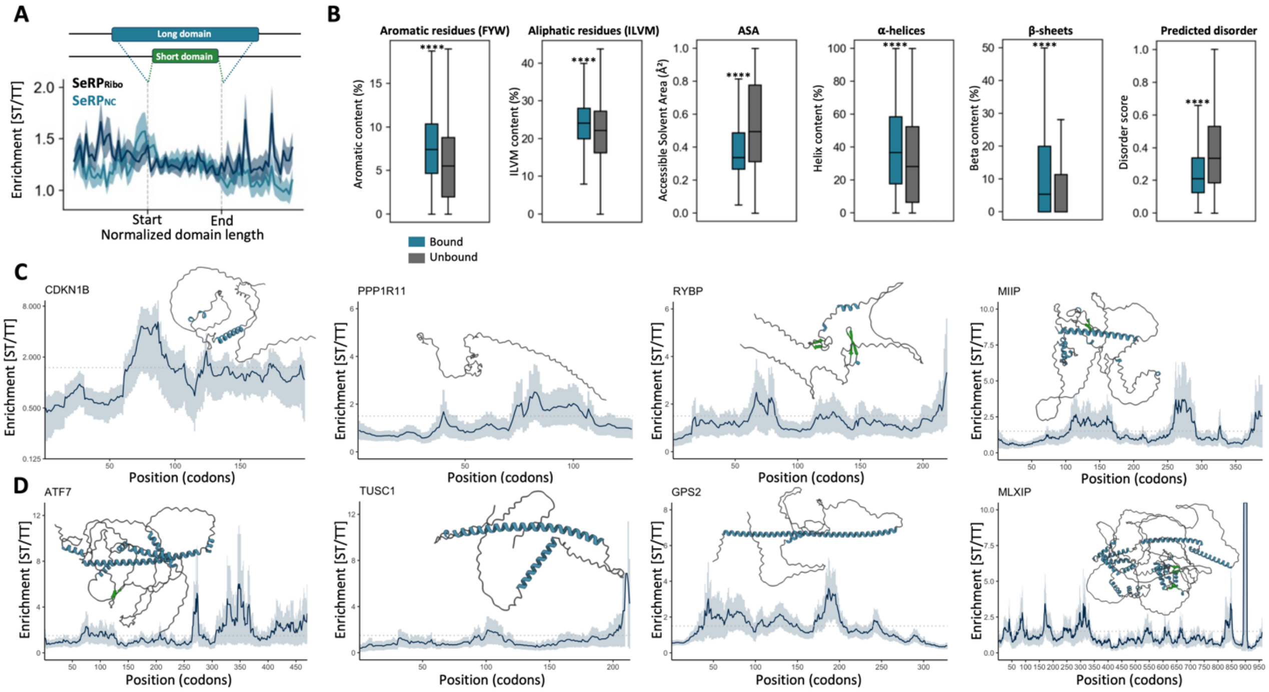

**Figure S5: Features and enrichment analysis of NAC-bound regions in ER-targeted, cytonuclear and non-globular proteins:** (**A**) Length-normalized ST over TT enrichment of ER-targeted proteins within domains and in flanking regions for NC-independent (black) and NC-dependent (blue) interactions. Shadowed areas indicate 95% confidence intervals. (**B**) Features of exposed regions bound and not bound by NAC based on the SeRP$_{NC}$ dataset. We consider regions bound by NAC to extend from the last (C-terminal) 60 residues synthetized at the onset of a binding period (30 residues in the exit tunnel and 30 residues exposed to the solvent) to the last residue emerging from the tunnel exit at the moment of NAC dissociation (end of binding period). To reduce overlaps, unbound regions are spaced 30 residues from the beginning or the end of a bound period. The first 30 codons of a protein sequence are not considered, since the polypeptides has not yet emerged from the exit tunnel. Statistical significance was tested using a two-sided Mann-Whitney U test. ****: p-value < 0.0001. (**C-D**) NC-dependent ST over TT enrichment of non-globular proteins (C) and proteins containing long coil-coils (D). Structures were extracted from the Alphafold database[1,2]. Shadowed areas indicate 95% confidence interval.

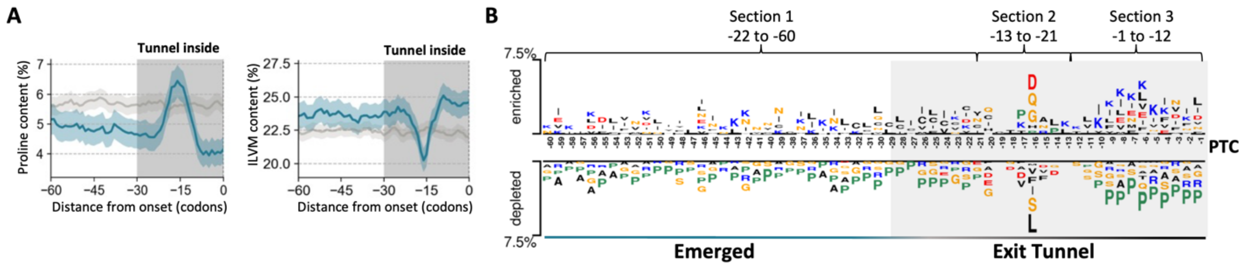


**Figure S6: Amino acid composition at the onset of NAC binding in cytonuclear proteins**. (**A**) Metagene profile of proline content and ILMV content aligned to the onset of NAC binding periods in cytonuclear proteins (blue) and in a randomized sequence control (grey). Dotted, dashed and solid lines represent sections 1, 2 and 3 in panel B, respectively. Shadowed areas indicate 95% confidence intervals. (**B**) Position-specific enrichment of residues in the ribosome-proximal 60 residues at the onset of NAC binding to nascent cytonuclear proteins.

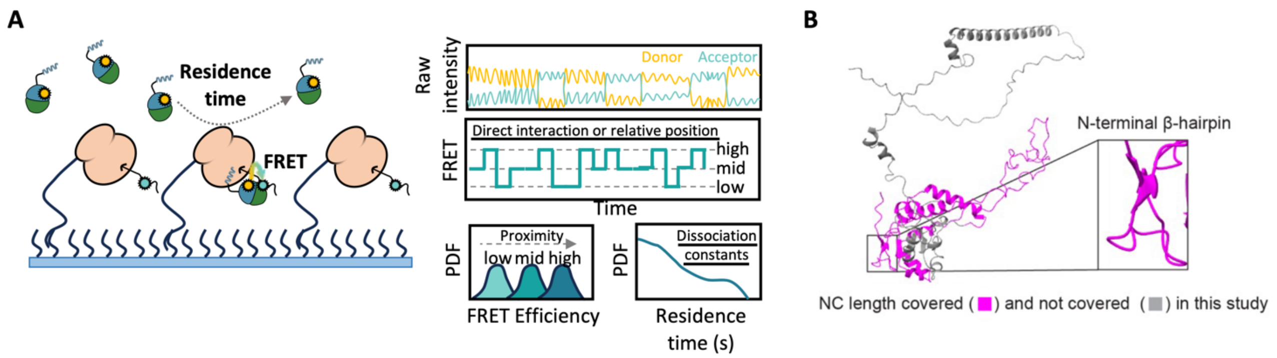


**Figure S7: Single molecule FRET set-up for studying NAC interaction with the ribosomes and the RPL4 nascent chains.** (**A**) Schematic figure showing the smFRET TIRFM approach and the results obtained with this strategy. (**B**) AlphaFold model of RPL4 with the experimentally tested NC region in *magenta* and the remainder of the protein in *grey*.

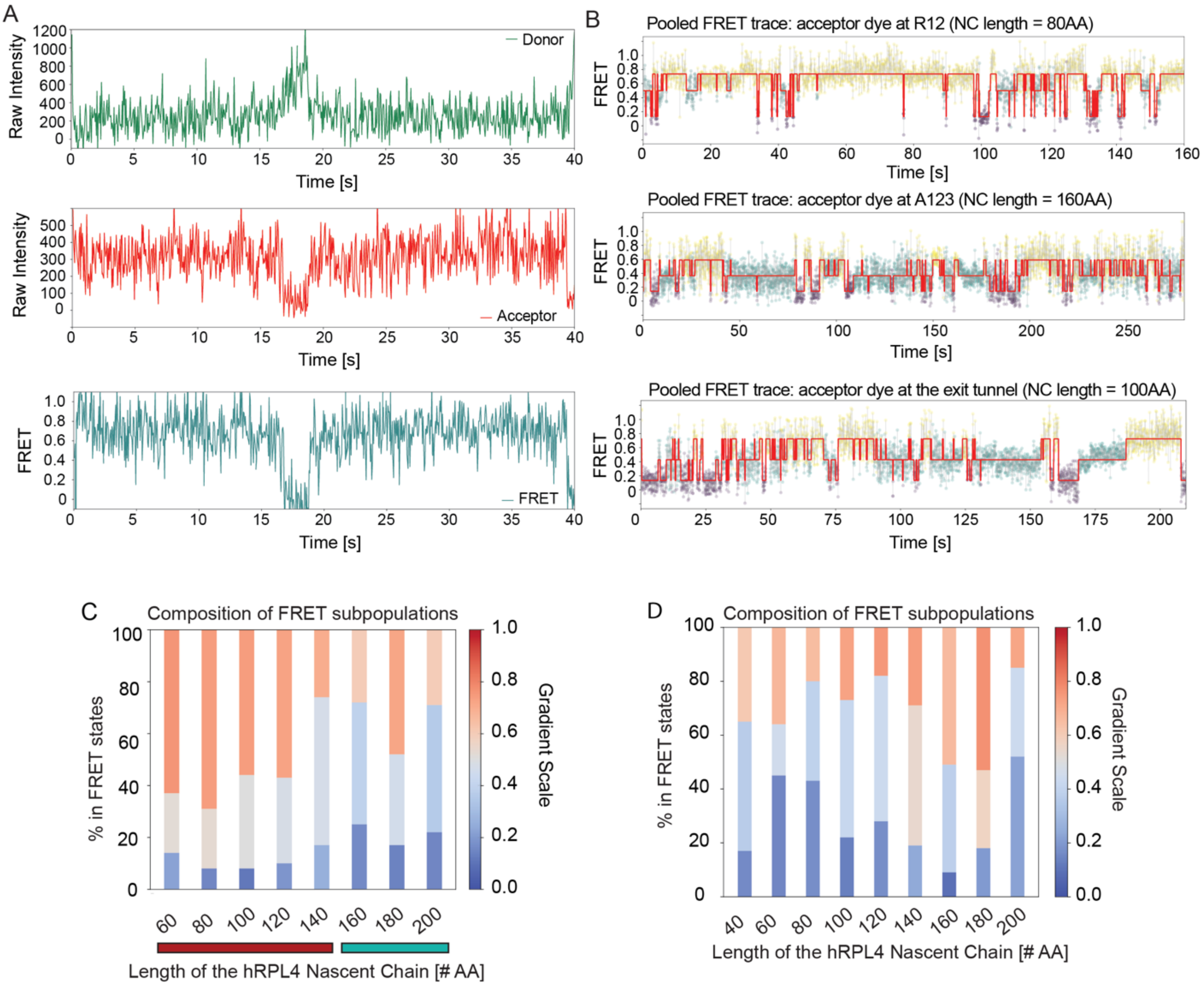


**Figure S8: Representative data and analyses of single molecule FRET traces.** (**A**) Representative traces showing raw intensity of the donor dye (top panel), acceptor dye (middle panel), and calculated FRET efficiencies (lower panel). (**B**) Hidden Markov modeling (HMM) were performed on pooled smFRET traces to establish the number of FRET states and the mean FRET efficiency of each state. A representative trace is shown for each FRET dye pair. Yellow, green, and purple color the region of the data assigned to the high, medium, and low FRET states. (**C, D**) Summary of the populations of NAC in different FRET states at the indicated NC lengths for the FRET pair with the acceptor dye at RPL4 residue 12 or 123 (C) or with the dye at the ribosome exit tunnel (D). The mean FRET efficiency of each state is color coded.

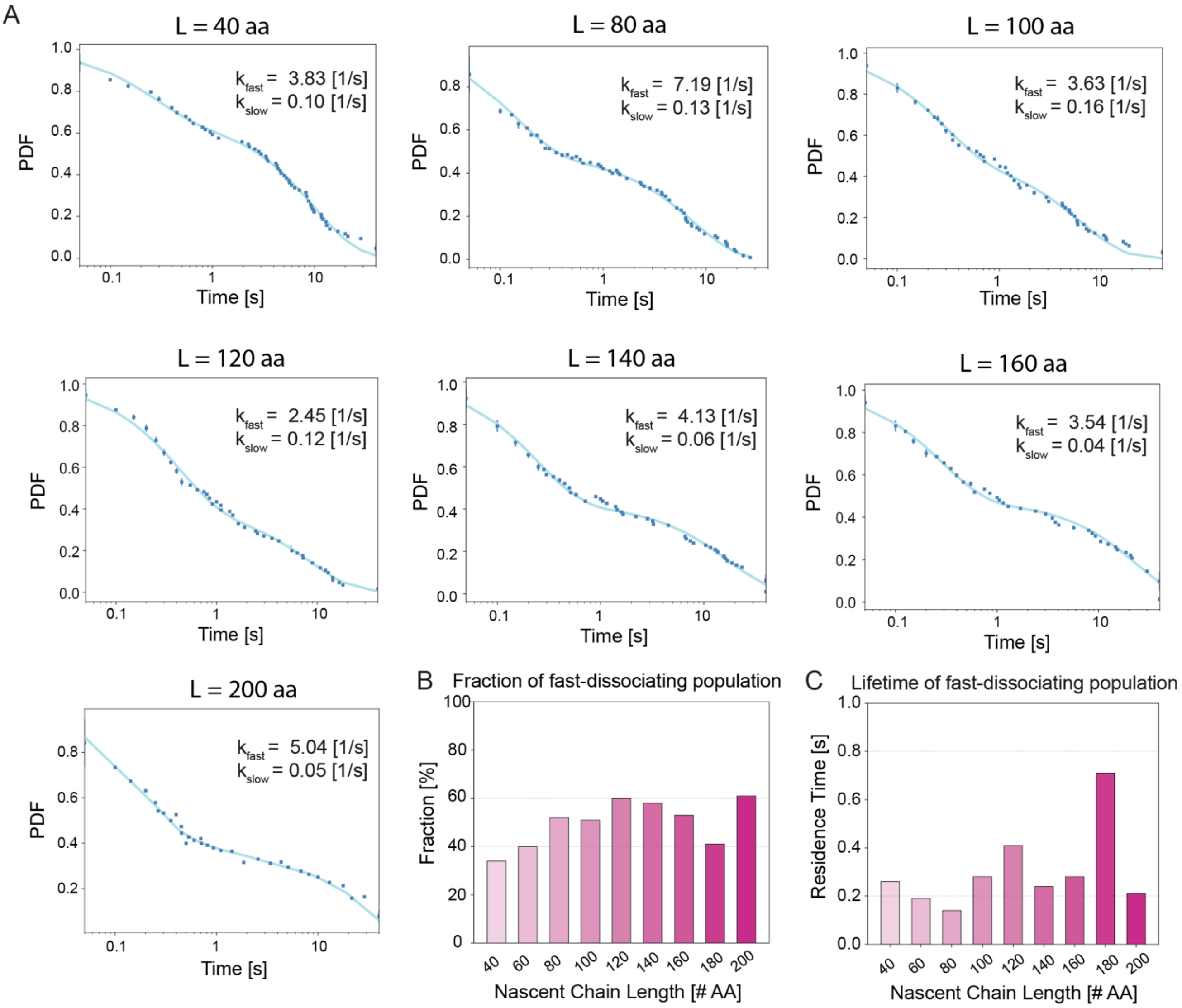


**Figure S9: Kinetic stability of NAC-RNC interaction.** (**A**) Time traces showing the dissociation of NAC from RNCs bearing the RPL4 NC at the indicated lengths. The dissociation rate constants of the two populations are indicated. (**B**) Summary of the % of NAC in the fast-dissociating population. (**C**) Summary of the residence time of the fast-dissociating population of NAC.

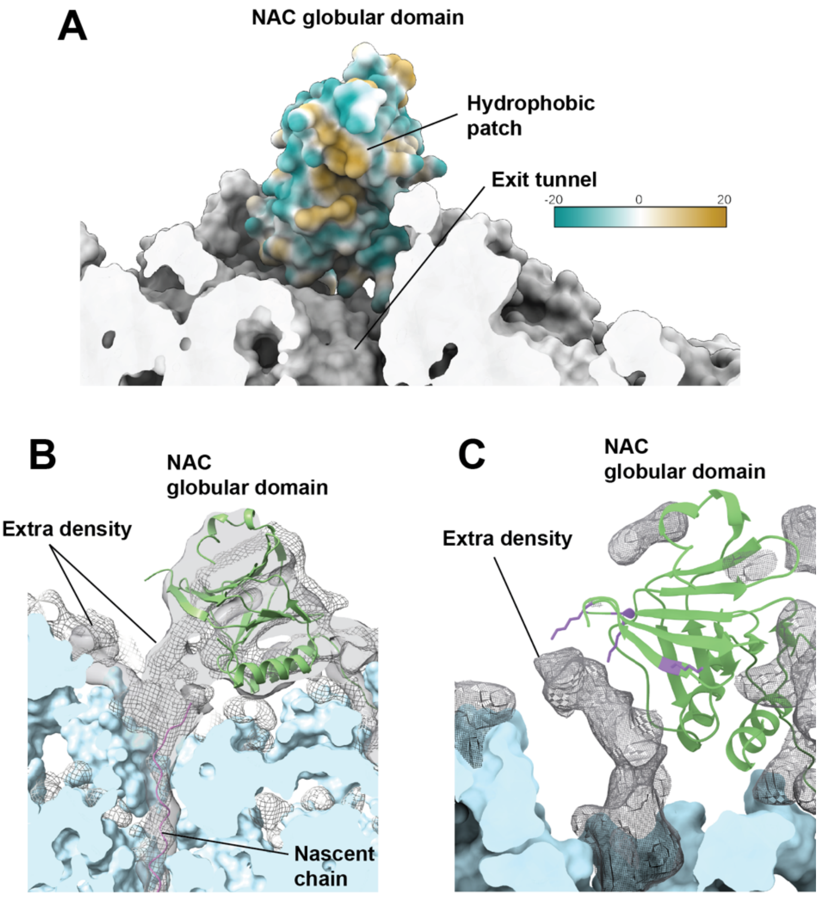


**Figure S10: Additional information on NAC and nascent chain density contacts.** (**A**) Molecular lipophilicity plot of the NAC globular domain (cyan – hydrophilic, yellow – hydrophobic). The hydrophobic patch facing the polypeptide exit tunnel is indicated. NAC is shown with respect to 60S (light gray) in the same perspective as in Figure 5E. (**B**) Comparison of the nascent chain density outside the tunnel exit in two independent datasets. The difference density between the refined cryo-EM maps and the 80S model, low-pass filtered to 6 Å, is shown in gray surface (dataset 1) and mesh (dataset 2). 60S proteins and rRNA is shown in light blue, NAC in green and the nascent chain in pink. (**C**) Lysines (violet) on the tunnel exit-facing surface of the NAC globular domain in proximity to the extra nascent chain density. The difference density between the cryo-EM map and the 80S/NAC model, low-pass filtered to 6 Å, is shown in black mesh. 60S proteins are shown in light blue and NAC in green.

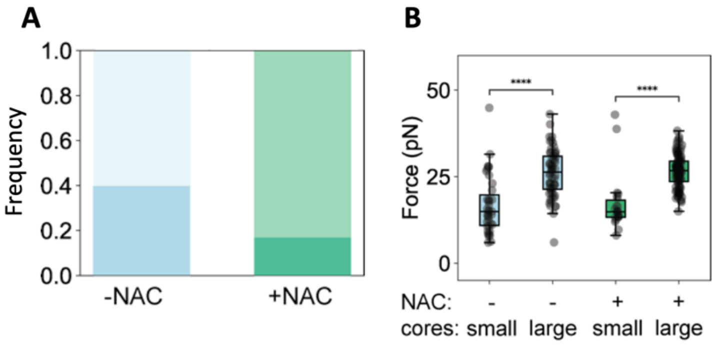


| NAC | - | - | + | + |
|---|---|---|---|---|
| Core | small | large | small | large |
| Force (pN) | 15 | 26 | 15 | 27 |

**Figure S11: NAC reduces the formation of non-native MBP cores.** (**A**) Frequency of small (in between 70 and 90 nm, dark color) and large (larger than 90 nm, light color) MBP cores in absence and presence of NAC. (**B**) Unfolding forces of small in between 70 and 90 nm ) and large (larger than 90 nm) MBP cores formed without (blue) and with (green) NAC (5 μM). **** indicate significant difference (P<0.0001; Mann-Whitney test). (**C**) Median values of small and large cores unfolding forces in absence and presence of NAC. Related to panel B.

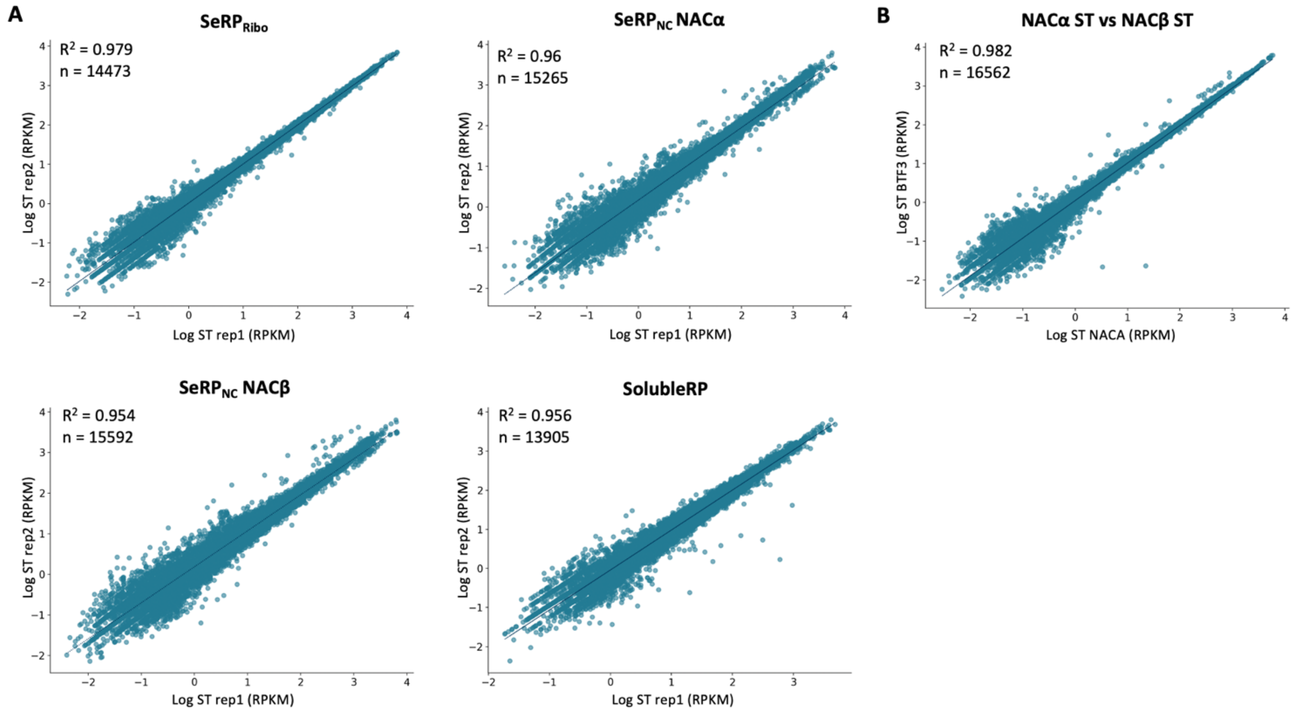


**Figure S12: Reproducibility between SeRP ST replicates** (**A**) Comparison of replicates of the ST of SeRP$_{Ribo}$, SeRP$_{NC}$ (immunoprecipitating NACα or NACβ) and SolubleRP ST. (**B**) Comparison of SeRP$_{NC}$ ST obtained by pulling on NACα or NACβ.

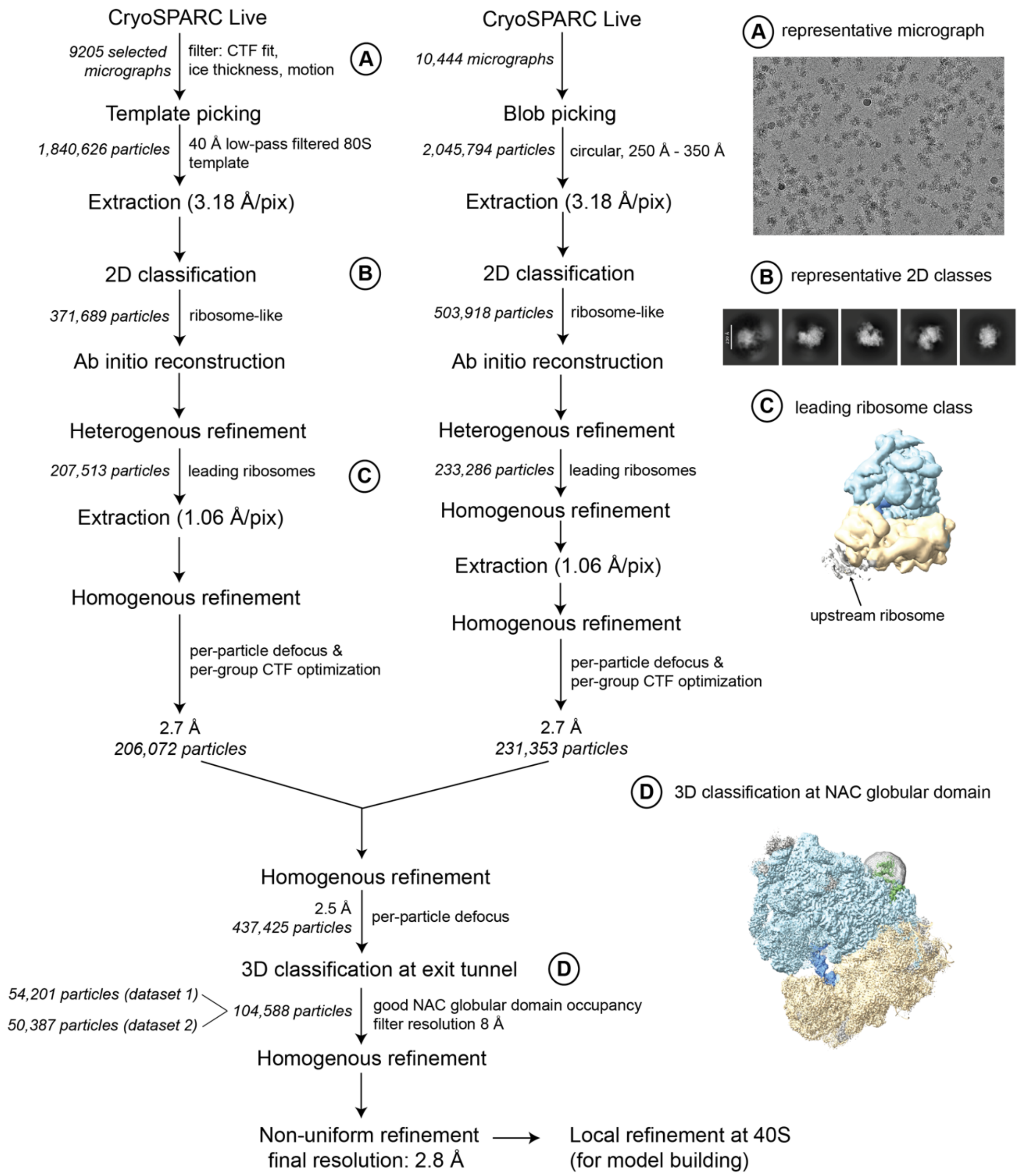


**Figure S13: Cryo-EM data processing flowchart for two cryo-EM datasets**. The initial processing of the two datasets was separate, shown in the two columns. 40S proteins and rRNA are colored in yellow, 60S proteins and rRNA in light blue, P-site tRNA in dark blue and NAC in green. The mask used for focused classification at the exit tunnel is represented as gray mesh.

**Table S1. Positive proteins and binding peaks per location.** Only proteins with RPKM values higher than 10 in the TT were included in the analysis.

| | **Cytonuclear** | **ER targeted** | **Mitochondrial** | **Overall** |
|---|---|---|---|---|
| Total | 4468 | 1198 | 441 | 7154 |
| NAC substrates | 3275 | 515 | 316 | 4584 |
| % of positive proteins | 73.3 | 43 | 71.6 | 64.1 |
| Positive binding periods | 12023 | 1087 | 782 | 15097 |

### Table S5: EM data collection and coordinate refinement statistics

| Complex | Human Xbp1u-stalled RPL4 RNC/NAC |
|---|---|
| EMDB code | EMD-##### |
| PDB code | #### |
| **Cryo-EM data collection and processing** | |
| Microscope | TFS Titan Krios G3i |
| Voltage (keV) | 300 |
| Camera | Gatan K3 |
| Magnification | 81,000 x (nominal) |
| Pixel size (Å) | 1.06 (super-resolution pixel at 0.53Å/pixel) |
| Electron exposure ($e^{-}$/Å$^2$) | 50 |
| Defocus range (µm) | (-0.7) – (-2.2) |
| Automation software | EPU |
| Energy filter slit width | 20 eV |
| Micrographs collected | 20,195 |
| Micrographs used | 19,649 |
| Initial particle images (no.) | 3,886,420 |
| Final particle images (no.) | 104,588 |
| Map resolution (Å) | 2.8 |
| Resolution range (local, Å) | 2.4 – 12 |
| Map sharpening B factor (Å$^2$) | -41.3 |
| **Coordinate real space refinement in PHENIX (version 1.21.2-5419)** | |
| Model resolution at FSC=0.5, masked (Å) | 3.0 |
| $CC_{mask}$ | 0.85 |
| Model composition | |
| Non-hydrogen atoms | 220,411 |
| Protein residues | 11,992 |
| RNA residues | 5,776 |
| Ligands: $Mg^{2+}$/$Zn^{2+}$/unassigned ions | 148/8/108 |
| B factors min/max/mean (Å$^2$) | |
| Protein | 33/483/186 |
| RNA | 32/754/201 |
| Ligand | 52/489/159 |
| **Model validation** | |
| *General* | |
| R.m.s. deviations | |
| Bond lengths (Å) | 0.002 |
| Bond angles (°) | 0.401 |
| MolProbity score | 1.1 |
| Clashscore | 3.5 |
| *Protein* | |
| Poor rotamers (%) | 0.8 |
| Cβ outliers (%) | 0.0 |
| CaBLAM outliers (%) | 1.3 |
| EM Ringer score | 2.7 |
| Ramachandran plot | |
| Favored (%) | 98.32 |
| Allowed (%) | 1.67 |
| Disallowed (%) | 0.01 |
| *RNA* | |
| Pucker outliers (%) | 0.5 |
| Bond outliers (%) | 0.09 |
| Angle outliers (%) | 0.0 |
| Suite outliers (%) | 13.4 |

**References**


1. Jumper, J., Evans, R., Pritzel, A., Green, T., Figurnov, M., Ronneberger, O., Tunyasuvunakool, K., Bates, R., Žídek, A., Potapenko, A., et al. (2021). Highly accurate protein structure prediction with AlphaFold. Nature *596*, 583–589. https://doi.org/10.1038/s41586-021-03819-2.

2. Varadi, M., Bertoni, D., Magana, P., Paramval, U., Pidruchna, I., Radhakrishnan, M., Tsenkov, M., Nair, S., Mirdita, M., Yeo, J., et al. (2024). AlphaFold Protein Structure Database in 2024: providing structure coverage for over 214 million protein sequences. Nucleic Acids Res *52*, D368–D375. https://doi.org/10.1093/nar/gkad1011.
